# Development of a whole-cell biosensor for ethylene oxide and ethylene

**DOI:** 10.1101/2024.02.19.581074

**Authors:** Claudia F. Moratti, Sui Nin Nicholas Yang, Colin Scott, Nicholas V. Coleman

## Abstract

Ethylene and ethylene oxide are widely used in the chemical industry, and ethylene is also important for its role in fruit ripening. Better sensing systems would assist risk management of these chemicals. Here, we characterise the ethylene regulatory system in *Mycobacterium* strain NBB4 and use these genetic parts to create a biosensor. The regulatory genes *etnR1* and *etnR2* and cognate promoter P_etn_ were combined with a fluorescent reporter gene (fuGFP) in a *Mycobacterium* shuttle vector to create plasmid pUS301-EtnR12P. Cultures of *M. smegmatis* mc^2^-155(pUS301-EtnR12P) gave a fluorescent signal in response to ethylene oxide with a detection limit of 0.2 µM (9 ppb). By combining the epoxide biosensor cells with another culture expressing the ethylene monooxygenase, the system was converted into an ethylene biosensor. The co-culture was capable of detecting ethylene emission from banana fruit. These are the first examples of whole-cell biosensors for epoxides or aliphatic alkenes. This work also resolves long-standing questions concerning the regulation of ethylene catabolism in bacteria.

## Introduction

Biosensors are devices that use biological components to detect or quantify a particular analyte; these may be deployed in a wide range of contexts including industrial facilities, environmental management, and healthcare ^1^. Biosensors contain two components - a biological sensor and a physical transducer – and are broadly classified based on the nature of each of these components ^2,3^. The biological components of the sensors include enzymes, proteins, antibodies, nucleic acids, phages or whole cells, while the physical transducers could be optical, electrochemical, thermoelectric, piezoelectric or electrochemical. Transcription-factor based biosensors are a subset of whole-cell biosensors that make use of regulatory proteins and their cognate promoters. In their original contexts, transcription factors modulate gene expression in response to an intracellular or environmental signal, while in biotechnology, they are instead often linked to expression of a fluorescent protein as the transducer ^4^. Transcription-factor based biosensors are particularly valuable in metabolic engineering and can be used to screen or select for desirable properties ^5,6^. They are cost-effective to produce and can be engineered to have high sensitivity and selectivity ^7^.

Aerobic bacteria that grow on alkanes or alkenes use monooxygenase (MO) enzymes as the first step in the oxidation of these substrates, yielding epoxides or alcohols, respectively ^8^. Bacterial MO’s are typically inducible by the initial hydrocarbon substrate or a downstream metabolite; this is a sensible strategy for the host cell given both the exotic nature of the substrates and the physiological stress of MO expression ^9^. In the alkane-oxidising bacteria, regulatory systems controlling MO expression are well-characterised, and many have been developed into biosensors for alkanes or their metabolites ^10–23^; this research field has been recently reviewed ^24^. In contrast, the regulation of alkene oxidation in bacteria has not been well-studied; some regulatory genes ^25–27^ and inducers ^28–30^ have been identified or predicted, but rigorous evidence for specific interactions between inducers, regulatory proteins and promoters has not been presented to date, limiting the development of this biotechnology.

Hydrocarbon biosensors have many potential applications, but to date, the primary focus has been on detection of alkanes (especially octane) as a marker for bioremediation of oil spills ^10,11,13,14,16,18,19,31^. Some of these alkane biosensors exploit the *alk* system, first intensively studied in *Pseudomonas putida* GPo1, which consists of the *alkBFGHJKL* structural genes and the *alkST* regulators ^10,32–34^. The AlkS regulator binds to C_6_-C_10_ alkanes, and thus overlaps with the substrate range of its cognate MO AlkB and acts as a direct biosensor for liquid alkanes. In other cases, these binding profiles are divergent; for example, the butane oxidation system of *Thauera butanivorans* involves induction of the BmoR regulator by 1-butanol ^35^, while the BmoXYBZDC MO preferentially attacks butane ^36^. Thus, although it controls growth on alkanes, this is an alcohol biosensor rather than an alkane biosensor ^23^.

Some Actinobacteria can grow on ethylene via a MO-mediated pathway that proceeds via ethylene oxide (epoxyethane) as the first intermediate ^26,27,37–39^. A subset of these bacteria also grow on the chlorinated derivatives of these compounds (vinyl chloride (VC) and chlorooxirane), and are of special interest for bioremediation. The initial ethylene/VC-oxidising enzyme EtnABCD (a soluble di-iron monooxygenase (SDIMO)), and the epoxide-transforming enzyme EtnE (epoxyalkane-coenzyme M transferase) have been partially characterised ^40,41^, however the downstream metabolic pathway and the regulation of this pathway are thus-far uncharacterised. The regulatory components of the ethylene-oxidation system are of great interest since these may provide the genetic parts for construction of an ethylene biosensor. Ethylene is not only the most important industrial organic chemical by volume ^42^, it is also the ripening hormone for many fruits and vegetables ^43^, and furthermore, it is a hazardous compound due to its high flammability ^44^. Therefore, an ethylene biosensor would be valuable, with multiple possible commercial applications.

Recent preliminary work identified two putative regulatory genes, *etnR1* and *etnR2*, associated with the ethylene oxidation genes in *Mycobacterium* strain NBB4, and showed via gel-shift assay that purified EtnR1 protein bound to a DNA region predicted to contain a promoter ^45^. Here, we aimed to confirm and extend these initial observations, by showing that EtnR1 and EtnR2 are indeed the regulators of the ethylene oxidation pathway, by localising their cognate promoter, and by investigating the inducer range and kinetic properties of the regulatory system. Finally, we aimed to use these genetic components to develop a prototype of a whole-cell ethylene biosensor.

## Results and Discussion

### Bioinformatic analysis of the *etnR1* and *etnR2* regulators and P_etn_ promoter

The predicted ethylene catabolism genes are located on a 144 kb plasmid in *Mycobacterium* NBB4 (Genbank CP003055). Inspection of this locus (Figure 1) reveals two possible regulatory gene candidates, *etnR1* (MYCCH_ RS29055) and *etnR2* (MYCCH_RS29050). Several lines of bioinformatic evidence support a role for both these genes in the regulation of the ethylene MO and other alkene catabolic genes, as described below.

**Figure 1.**
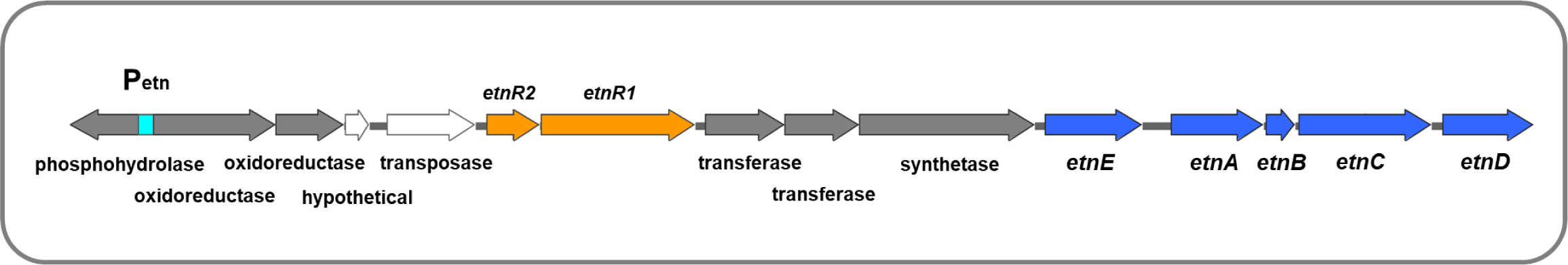
Ethylene metabolic genes and regulators in strain NBB4. Metabolic genes of known function are in blue, predicted metabolic genes are in grey, and predicted regulatory genes are in orange. The predicted promoter region (P_etn_) is in cyan. The region shown is 16.5 kb in size, and represents only part of the complete ethylene catabolism and coenzyme M biosynthesis locus, which is approx. 40 kb.

The *etnR1* and *etnR2* genes are embedded within a larger gene cluster containing the *etnABCD* MO and several other genes either known or suspected to play a role in alkene metabolism ^26^; these are all in the same orientation, suggesting a common promoter, i.e. they form an operon. It is typical for transcription activators to have a low level of constitutive expression that is upregulated in the presence of the correct inducer ^46^; thus we postulate that EtnR1 and EtnR2 are initially expressed at low levels from an as-yet-unidentified constitutive promoter immediately upstream, and then are upregulated via a positive feedback loop upon activation of the operon in the presence of the hydrocarbon inducer. There is precedent for this in the AlkS system of *P. oleovorans* ^47^.

BLAST analysis of EtnR1 and EtnR2 provides evidence that these comprise a two-component sensing system, with EtnR2 as the sensor, and EtnR1 as the DNA-binding protein. EtnR1 is a member of the CdaR superfamily of regulatory proteins (COG3835), with homology to CdaR over the region from amino acids (aa) 250-554 (E-value = 1.8e^-21^). The CdaR regulator itself is an activator of sugar diacid catabolism in *E. coli* ^48^. A helix-turn-helix (HTH) domain, common to DNA-binding proteins, is present at the C-terminus of EtnR1 (aa 501-561), with homology to the HTH domain of the *Bacillus subtilis* PucR regulator; this regulator controls purine catabolism in *B. subtilis* ^49^. EtnR1 also contains a GAF domain (aa 135-285) similar to those found in transcriptional activators such as FhlA and NifA; GAF domains are common in sensory proteins^50^.

EtnR2 contains just one conserved domain that spans most of the protein (aa 21-174, E value = 3.42e-46); this is a <u>me</u>thanogen/methylotroph <u>D</u>cmR <u>s</u>ensory (MEDS) domain. MEDS domains are involved in the sensing of small hydrocarbons and their derivatives and were first identified in methanogens and methylotrophs ^51^. EtnR2 has 24% aa identity and 34% aa similarity to the archetypal protein in this family, the DcmR dichloromethane sensor of Methylobacterium sp, strain DM4 ^52^. EtnR2 lacks the N-terminal HTH domain seen in DcmR; this is unsurprising since DcmR acts as a single stand-alone regulator (transcriptional repressor) while EtnR2 most likely acts together with EtnR1. We hypothesise that ethylene or a metabolite derived from ethylene binds to EtnR2, which transfers this signal to EtnR1, which then binds to the P_etn_ promoter region, leading to activation of expression of the ethylene catabolic genes.

Homologs of *etnR1* and *etnR2* are present within the predicted ethylene catabolic gene clusters of all other known ethylene/vinyl chloride-oxidising Actinobacteria (Figure 2), and these homologs are highly conserved in sequence and location relative to the genes in strain NBB4 (Genbank NC_018023). However, more distant homologs also occur in Actinobacteria with no known capability for hydrocarbon oxidation; these can be subdivided into one group which co-occur with genes similar to those involved in alkene oxidation and/or CoM biosynthesis (these may be bacteria with yet-to-be-demonstrated alkene oxidation capacities), and another group which do not colocalise with catabolic genes. The EtnR1/R2 regulatory system appears to be a specialised subset of a broader group of regulators with thus-far unknown functions, which are common in the Actinobacteria.

**Figure 2.**
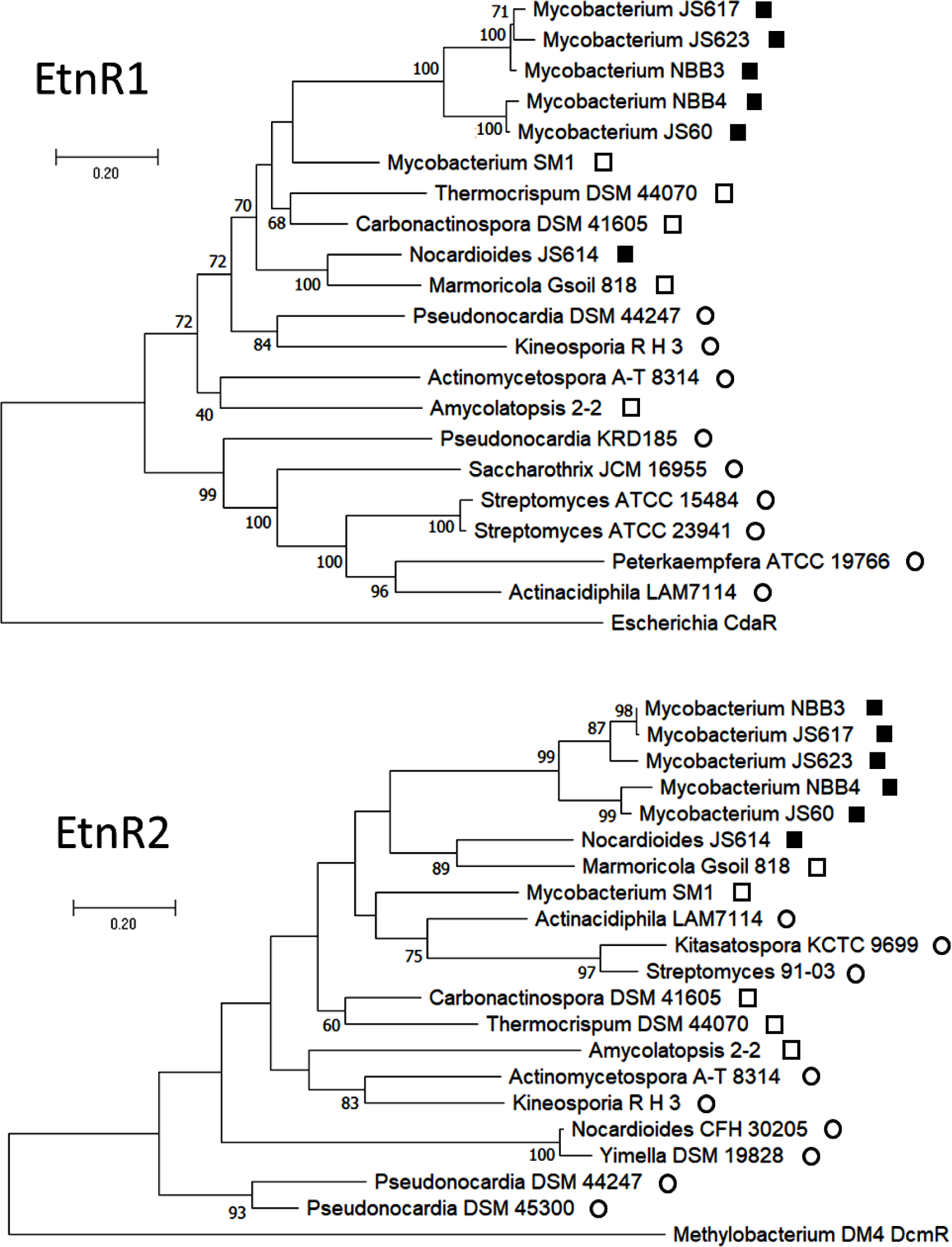
Phylogeny of EtnR1 and EtnR2 homologues. EtnR1 or EtnR2 homologues were detected by BLASTp, and representative sequences with 40-99% identity to the NBB4 proteins selected for analysis, along with one representative outgroup (CdaR or DcmR). Alignments were generated in MEGA using the MUSCLE algorithm, then trimmed (573 aa or 194 aa for EtnR1 and EtnR2, respectively). Trees were generated in MEGA11 by maximum likelihood method with 100 replicates. Symbols are as follows; 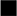, known alkene utilizers; 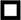, alkene catabolism genes near regulator; 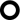, other types of genes near regulator. Bootstrap values >50% are shown at nodes.

Based on inspection of the NBB4 genome sequence, the promoter for the ethylene catabolic genes was predicted to occur in a 167 bp non-coding region that is 4 kb upstream of the regulatory genes (Figure 1). This region is positioned between two divergently-oriented gene clusters, with the predicted coenzyme M (CoM) biosynthesis genes on one side, and the ethylene catabolic genes on the other side. Note that coenzyme M is a key cofactor required for epoxide metabolism by the EtnE enzyme in the alkene catabolic pathway, thus it is logical the CoM biosynthetic gene clusters to be co-ordinately regulated with the alkene catabolic genes; in aerobic bacteria, the only known function of CoM to date is in alkene catabolism ^53^. We therefore predict that this non-coding region contains two divergently-oriented promoters controlled by the same regulators (i.e. EtnR1/EtnR2). Here we focus our attention just on one component of this region, i.e. the promoter driving the ethylene catabolic genes, which we have named P_etn_.

The predicted P_etn_ promoter (Figure 3) contains putative −35 and −10 sequences (TTGTCT, AATAAT) that are similar to the consensus for σ70 promoters in mycobacteria (TTGACR, TATRMT, and although the spacing of these elements is only 9 bp, this shorter spacing is not uncommon in mycobacteria ^54^. Motifs similar to the −24 and −12 σ^54^ consensus sequences (MRNRYTGGCACG, TTGCWNNW ^55^) are also identifiable in this region, although it should be noted that σ^54^ homologues (RpoN) have not been identified to date in mycobacteria ^56,57^ and there are no RpoN homologues detectable in the NBB4 genome. The putative P_etn_ promoter region contains several direct and inverted repeat sequences; these are typical of binding sites for transcription factors ^54^. Sequences similar to the binding sites of EtnR1 homologs such as PucR (WWWCNTTGGTTAA ^49^) or FhlA (CATTTCGTACGAAATG (Leonhartsberger et al., 2000)) could not be detected.

**Figure 3.**
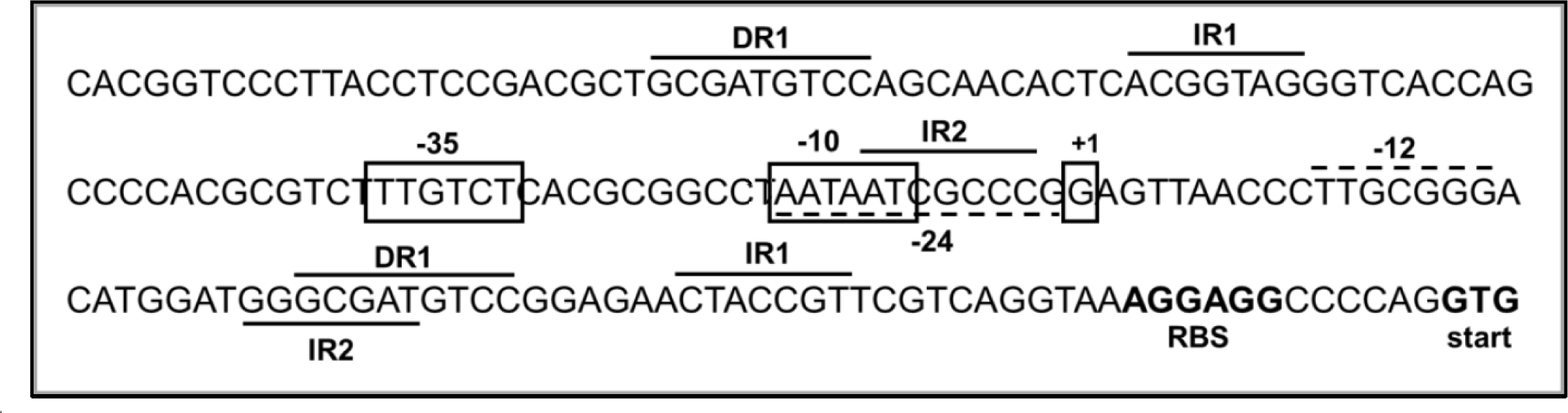
DNA sequence and features of predicted promoter region P_etn_. Predicted −35 and −10 sequences and transcription start (+1) are boxed. Predicted −24 and −12 sequences have dotted underlines or overlines. Inverted repeats (IR1, IR2) and direct repeats (DR1) have solid underlines or overlines. The predicted RBS and start codon for the oxidoreductase gene MYCCH_RS29030 are shown in bold.

### Design and construction of biosensor

*Mycobacterium smegmatis* mc^2^-155 was previously shown to be a good expression host for the ethylene MO EtnABCD ^40^ and thus we reasoned that mc^2^-155 would also be appropriate for functional expression of the EtnR1 and EtnR2 regulators. We first constructed a *Mycobacterium/E.coli* shuttle vector, pUS301, which was then used as the backbone for biosensor construction. The annotated sequence of pUS301 is provided in the Supporting Information; this vector contains the pUC origin for replication in *E. coli*, the pAL5000 origin for replication in *Mycobacterium*, a kanamycin resistance gene functional in both genera, and a cumate-inducible promoter for expression of cloned genes ^59^. The latter was created by fusing the CymR repressor binding sites (cumate operators; CuO) on each side of the strong P_C_ promoter from the class 1 integron ^60^.

The prototype biosensor was constructed via Golden Gate assembly of three synthetic DNA fragments (*etnR1*, *etnR2*, P_etn_) into pUS301, such that *etnR1* and *etnR2* were co-expressed from the cumate-inducible promoter (Figure 4). Both *etnR1* and *etnR2* were modified from the wild-type by removing internal restriction sites via single base changes and by reducing the overall GC content to enable gene synthesis. Both restriction site removal and changes to GC content were done while preserving the amino acid sequence using a manual codon harmonisation strategy, i.e. choices of codons in the modified sequence were matched as closely as possible to the relative frequency of the codon in the wild-type gene at that position. The 5’ ends of both regulatory genes were fused to the same synthetic strong ribosome binding site (AAGGAG), with a 6-bp spacer region before the start codons.

**Figure 4.**
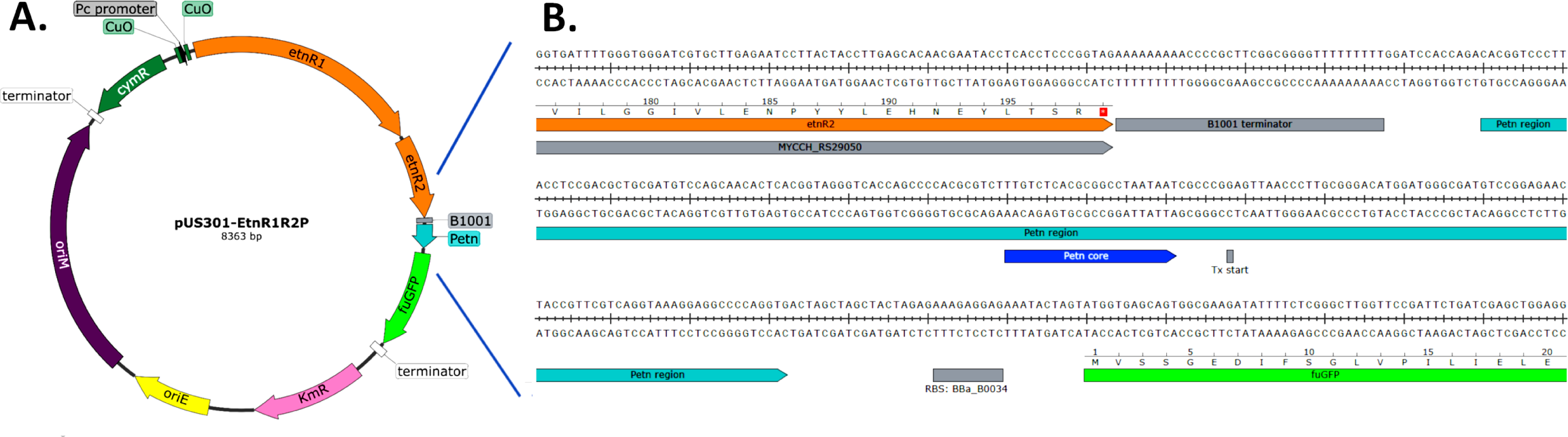
Schematic maps of biosensor plasmid pUS301-EtnR12P. Panel A. Overall plasmid map. Key features include: KmR (pink), oriE (yellow), pAL5000 oriM (purple), CymR (dark green). EtnR1 and EtnR2 (orange) and P_etn_ (turquoise). Map was constructed with SnapGene software. Panel B. Sequence details of transcription and translation signals controlling fuGFP in the biosensor plasmid.

To minimise transcriptional read-through of the cumate-inducible promoter into the P_etn_ promoter and its reporter gene, the strong bidirectional transcription terminator BBa_B1001 from the iGEM Parts Kit was added between *etnR2* and P_etn_. As a reporter gene to measure transcription, the gene for free-use green fluorescent protein (fuGFP; BBa_K3814004) was positioned immediately downstream of P_etn_, with 112-bp spacing between the predicted transcription and translation start points. The fuGFP gene was used with the strong RBS BBa_B0034 ^61^ separated from the start codon by a 7 bp spacer.

### Identification of inducers of the EtnR1-EtnR2-P_etn_ system

Cultures of mc^2^-155(pUS301-EtnR12P) were treated with cumate to induce the regulator genes, and various test compounds (alkenes, epoxides, alcohols) as possible inducers of the EtnR1-EtnR2-P_etn_ system (Figure 5). Cultures induced with cumate only gave the same low fluorescence as the solvent control (DMSO) (*p* >0.05), providing confidence that any increase in fluorescence would be due to P_etn_ being activated, and not read-through from the cumate-inducible promoter. A strong response was seen after ethylene oxide addition (EtO, 1 µM), with an average fluorescence 8.5-fold greater than the solvent control (*p* < 0.05). Weak induction (1.8-fold) was seen with ethylene glycol (EtGL, 1 µM) but this was not statistically significant (*p* = 0.9518 vs. uninduced). Ethylene (10% v/v in headspace) did not give significant fluorescence (*P* > .9999 compared to uninduced sample). Biosensor plasmid variants containing only *etnR1* or *etnR2* (pUS301-EtnR1P, pUS301-EtnR2P) produced no significant fluorescence after exposure to EtO, confirming that both genes are necessary for the system to function; this experiment also confirms that EtnR1 is an activator rather than a repressor, since if the latter were true, P_etn_ should have switched on when *etnR1* was removed, regardless of the presence of the epoxide inducer.

**Figure 5.**
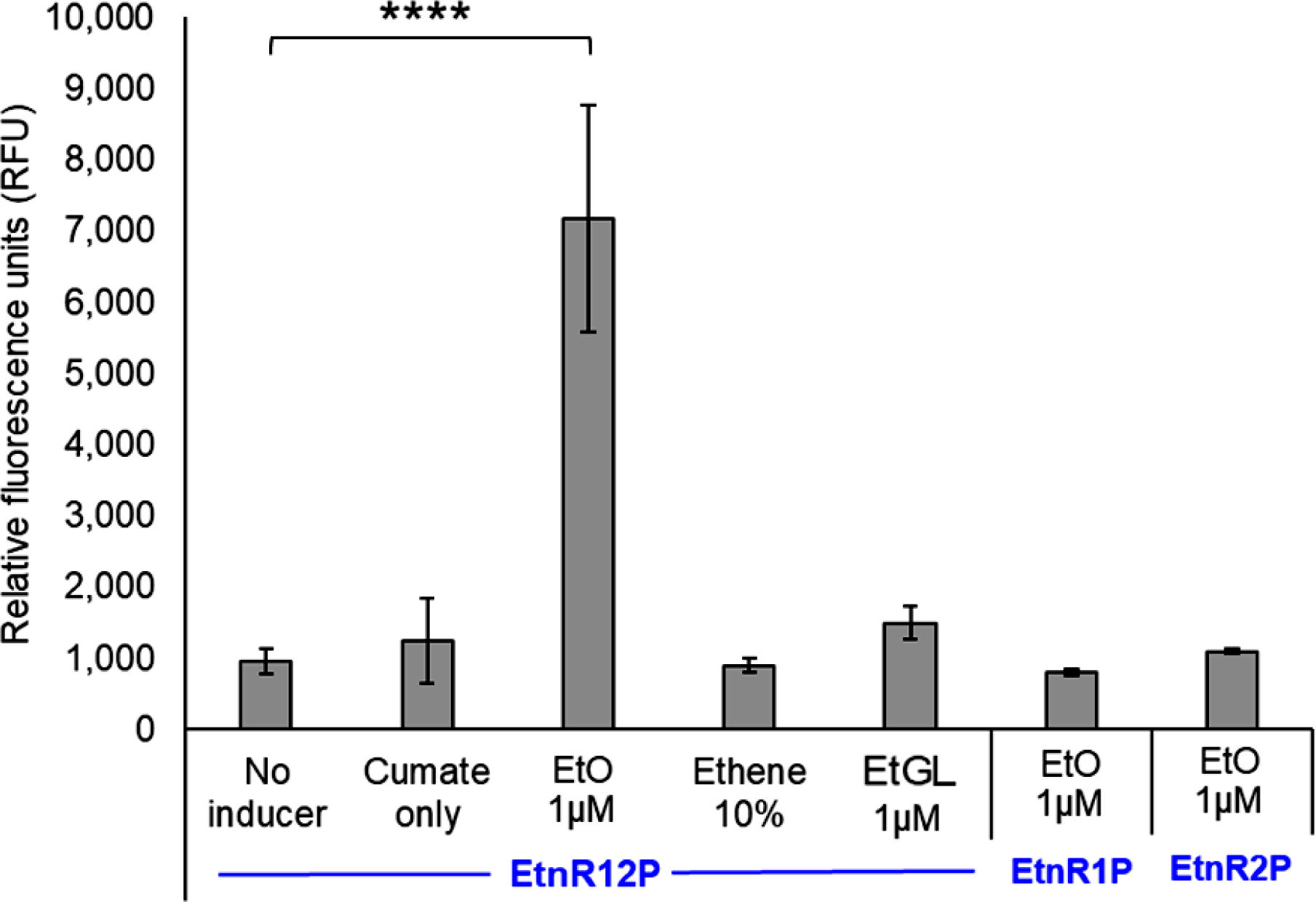
Testing inducers of the EtnR1-EtnR2-P_etn_ regulatory system. Liquid inducers (EtO, EtGL) were added as solutions in DMSO, while gases (ethylene) were added neat to the headspace. RFU values were derived by dividing the raw fluorescence readings by the density (OD_600_) of the cultures. Data were collected 16 h after inducer addition. Data show the mean of three biological replicates, and error bars show the standard deviation. Statistical analysis was carried out using one-way ANOVA and Tukey post-hoc multiple comparisons analysis, with p < 0.05 judged as ‘significant’. * P ≤ 0.05, ** P ≤ 0.01, *** P ≤ 0.001, **** P ≤ 0.0001.

### Characterisation of biosensor sensitivity and response range with EtO

The responses of mc^2^-155(pUS301-EtnR12P) cultures were tested with a range of EtO concentrations (Figure 6A). The minimum concentration giving a significant fluorescent response (i.e. the measured detection limit of the biosensor) was 0.2 µM EtO (= 9 ppb), and the response reached a maximum around 50 µM (= 2,200 ppb). It should be noted that these are nominal concentrations, i.e. assuming that all EtO is dissolved in the liquid phase; this is a reasonable approximation given the low Henry’s constant for ethylene oxide (*H^cp^*=5.8 x 10^-2^) ^62^. Calculation of the predicted amount of EtO in the liquid phase using the Henry’s constant gives a similar number (8.17 ppb; see Section S1 of the Supporting Information). The transfer function of the biosensor relating the inducer concentration (input) to the fluorescent signal (output) was modelled using the Hill equation, which is often applied to transcription factor-promoter interactions during the regulation of gene expression ^63,64^. The resulting sigmoidal curve when plotted on a logarithmic x-axis had a good fit (R^2^=0.9486) and allowed the derivation of key performance parameters (Figure 6B).

**Figure 6.**
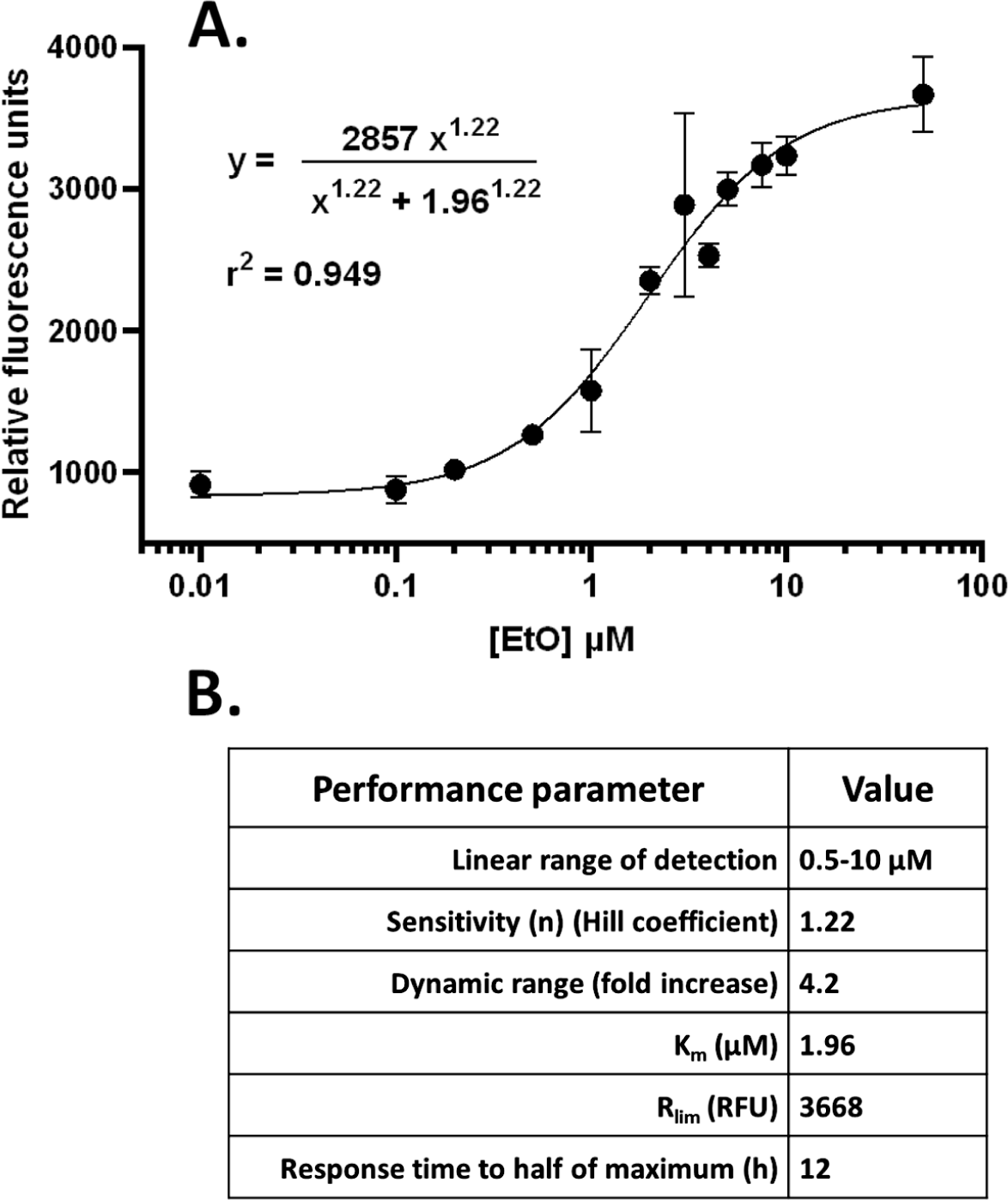
Epoxide biosensor response to EtO and performance parameters. Panel A. Plot of biosensor response at different EtO concentrations. RFU values were measured 10 hours after simultaneous induction with cumate and EtO. Data points are the mean of four biological replicates, and error bars show the standard deviation. The data have been fitted with a sigmoidal function via the equation shown on the graph. Panel B. Summary of performance parameters of biosensor with EtO.

### Time course of biosensor induction and effect of pre-induction of regulators

A time course assay was done to assess the induction profile of the biosensor with EtO (Figure 7), using cumate only (negative control), co-induction (cumate and EtO added simultaneously) and pre-induction (cumate added 16 h before EtO); the latter test aimed to determine if biosensor response could be accelerated by having the regulator proteins present in the cells before EtO was added. Pre-induction decreased the lag time before RFU began to increase by 1 h and gave a faster initial rate of increase in RFU (252 vs. 192 RFU/h; Figure 10-B and -C). Co-induction gave an initial linear increase in RFU over the first 18 h (r^2^=0.9959), while for pre-induction, the initial linear increase in RFU was only for 6.5 h (r^2^=0.9946), then the rate slowed to 115 RFU/hr between 6.5 – 22 h (R^2^ = 0.985). At 15 h post-induction, the RFU of the co-induced culture caught up with the pre-induced culture, and after this point, the cultures behaved similarly, reaching a peak RFU at 23 h, then declining. EtO induction did not dramatically impact the growth of cultures (Figure S1, Supporting information), although pre-induction led to a slowing of growth after 12 hours relative to co-induction. Overall, the pre-induction method was confirmed to give a more rapid biosensor response.

**Figure 7.**
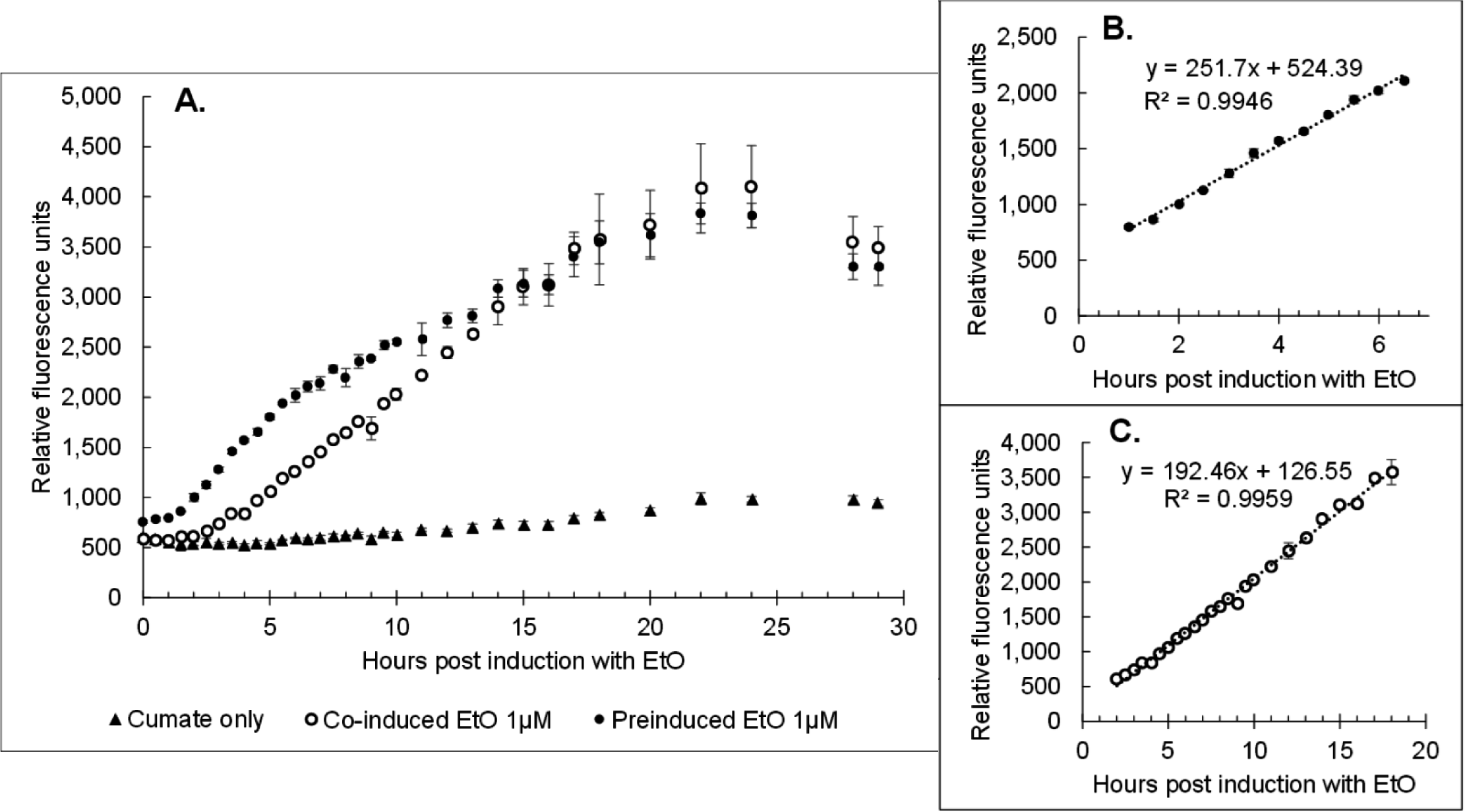
Time course of induction of EtO biosensor. A. Complete time course data of all sample sets. B. Linear fit through data from pre-induced data set between 1-7 hours post-induction with 1uM EtO. C. Linear fit through data from co-induced data set between 1-16 hours post-induction with 1uM EtO. Data are the means of three biological replicates, and error bars show the standard deviation

**Figure 8.**
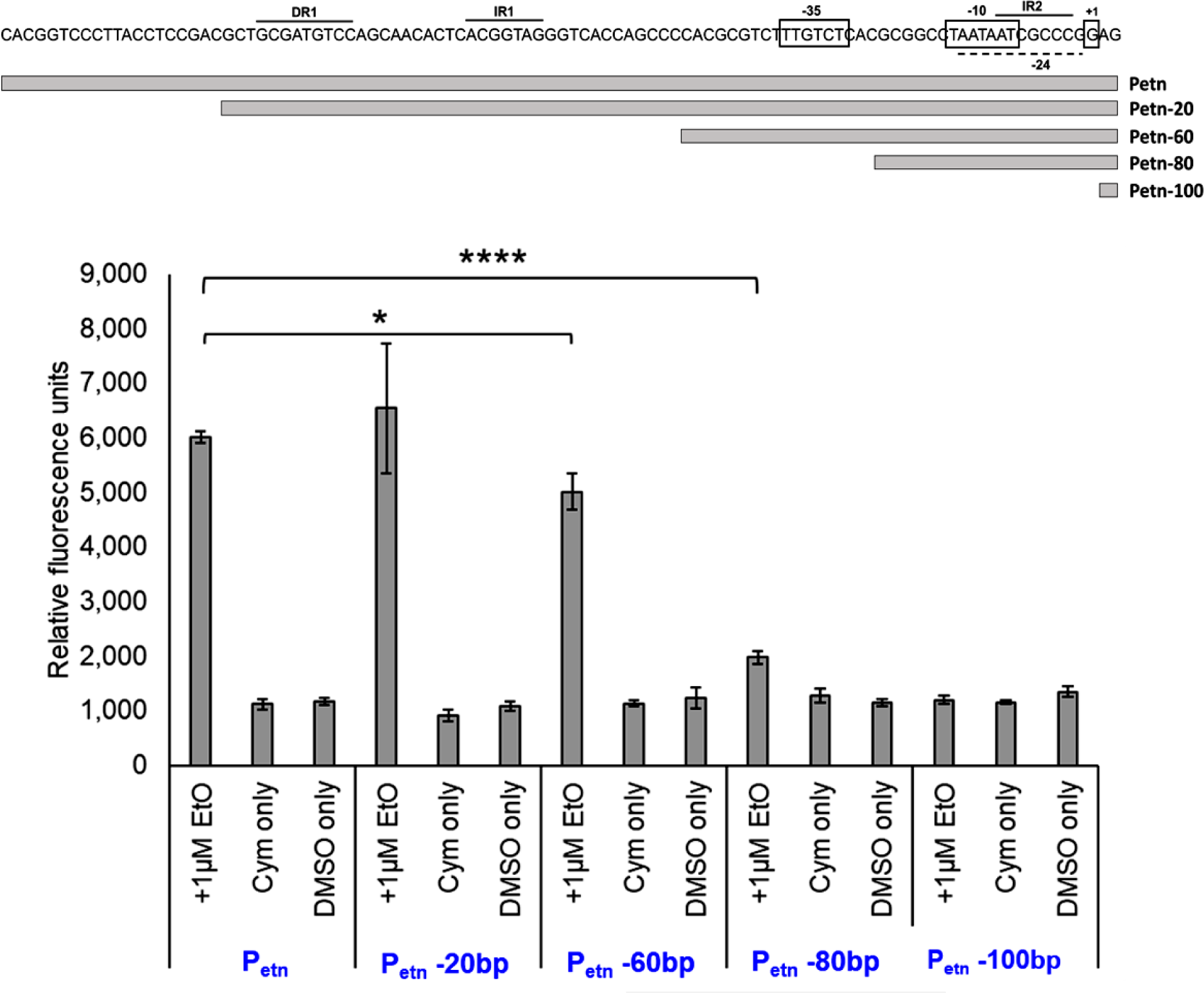
Localisation of P_etn_ promoter elements via 5’ sequence truncations. Fluorescence data was collected 21 h after induction with 1 µM EtO. Negative controls were induced with cumate only or cumate + DMSO. Data are means of four biological replicates, and error bars show the standard deviation. Statistical analysis was performed by two-way ANOVA, and p-values reported in the text are from Tukey post-hoc multiple comparisons analysis with p < 0.05 judged as ‘significant’. * P ≤ 0.05, ** P ≤ 0.01, *** P ≤ 0.001, **** P ≤ 0.0001.

**Figure 9.**
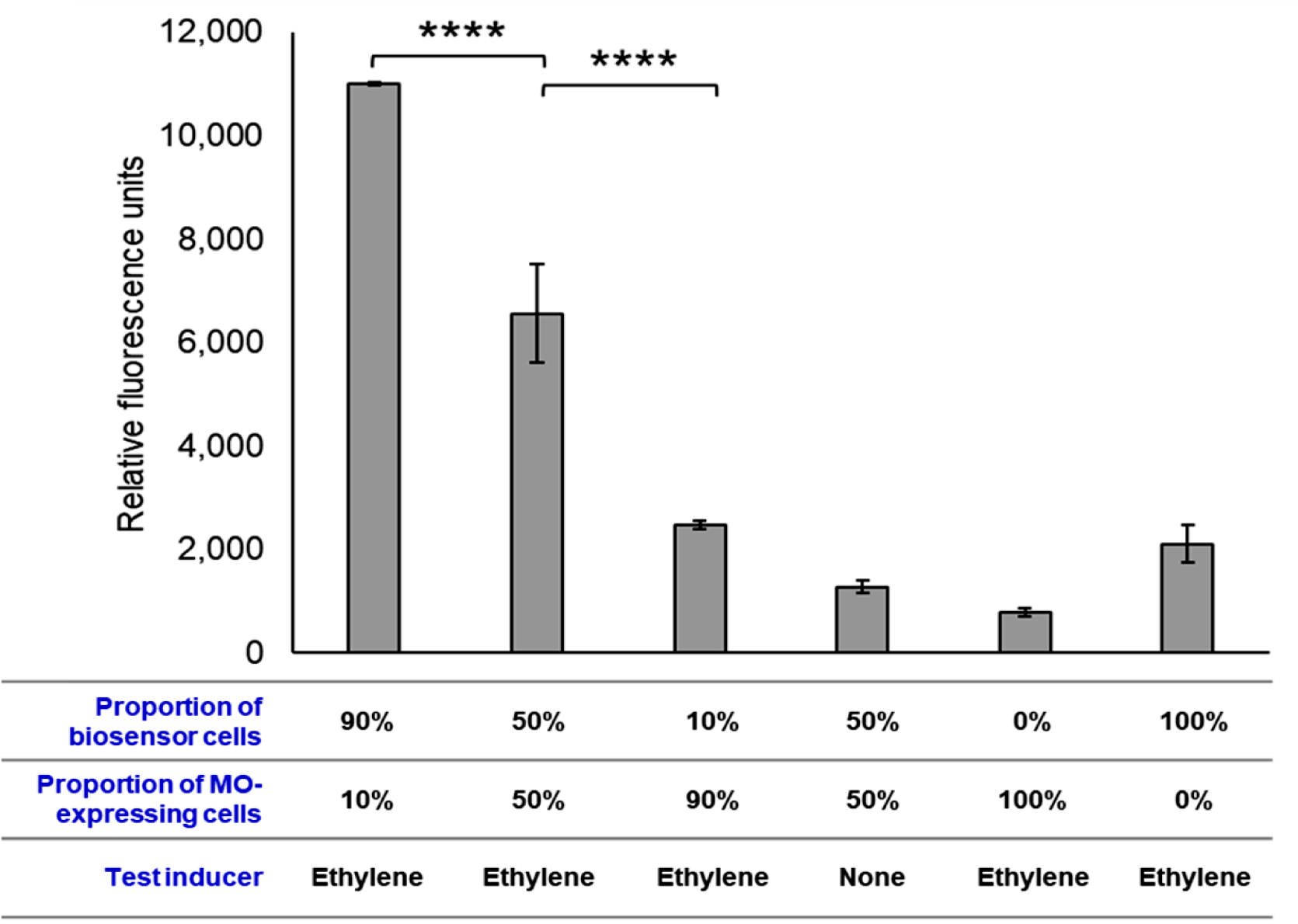
Ethylene detection by co-culture of EtO biosensor with MO-expressing cells. Cells of mc^2^-155(pUS116-etnABCD) were indcued with 1% acetamide, while cells of mc^2^-155(pUS301-EtnR12P) were induced with 100 µM cumate. RFU was calculated 21 hrs after ethylene addition. Data are the mean of three biological replicates. Error bars show the standard deviation. Statistical analysis was performed using one-way ANOVA, and p-values from Tukey post-hoc multiple comparisons analysis are presented with p < 0.05 judged as ‘significant’. * P ≤ 0.05, ** P ≤ 0.01, *** P ≤ 0.001, **** P ≤ 0.0001.

**Figure 10.**
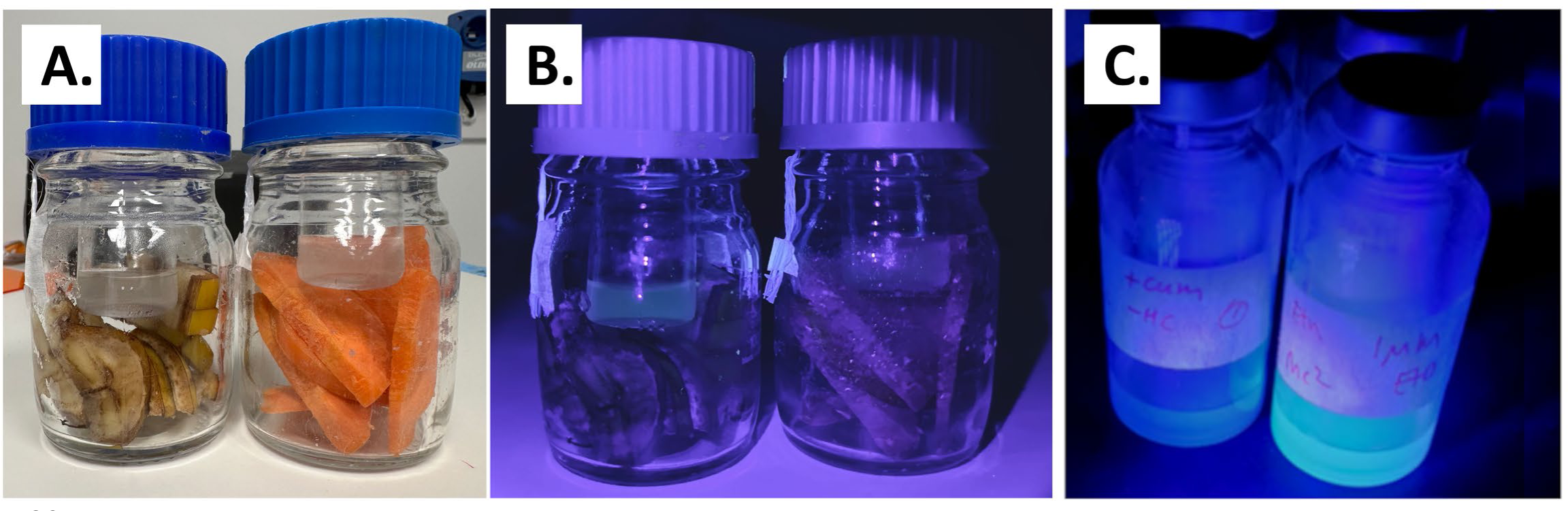
Detection of ethylene emission from banana by co-culture biosensor. Panel A. Visualisation of the experimental system after 3 days incubation under regular white light illumination. Panel B. Visualisation of the experimental system after 3 days incubation under long-wave UV light illumination. Panel C. Example of fluorescent response of biosensor to pure EtO (1 µM) under long-wave UV light illumination (these bottles are taken from the experiment shown in Figure 6).

### Localization of P_etn_ promoter sequence

To identify the minimum necessary promoter region, four variants of the 167 bp P_etn_ sequence with different 5’ truncations were assayed for activity (Figure 8). Removal of the first 20 bp of the sequence had no significant impact on biosensor response (p = 0.8381). A 60 bp deletion including one copy each of the DR1 and IR1 repeat elements gave a 20% decrease in RFU (p = 0.0377). The 80 bp deletion removed the putative −35 site and resulted in an 80% decrease in RFU; the signal from this construct was not significantly different from the 100 bp deletion construct after EtO induction (p = 0.2619) or the negative controls (p = 0.2040 no truncation uninduced, p = 0.1397 no truncation induced with cumate alone). The 100 bp deletion removed the putative −10 site and led to a complete loss of biosensor activity. Taken together, these results suggest that the P_etn_ promoter is of the σ^70^ type, that the −35 and −10 sequences identified here are the key sites for facilitating RNA polymerase binding, and that the DR1 and/or IR1 sequences may be involved in the binding of EtnR1 to the promoter region.

### Adding ethylene MO expands the biosensor range to include alkenes

Since alkenes can be converted into epoxides via the action of MO enzymes, we hypothesized that addition of a MO activity into our biosensor system would enable it to detect alkenes in addition to epoxides. Proof-of-principle for this approach was obtained via a co-culture method, via combining mc^2^-155 cells containing pUS301-EtnR12P with mc^2^-155 cells containing pUS116-etnABCD; the latter plasmid contains the ethylene MO of *Mycobacterium* NBB4 under the control of an acetamide-inducible promoter ^40^. Different initial ratios of the two cultures (50:50, 90:10, 10:90) were tested. A strong fluorescent response (equal or better than with ethylene oxide) was seen in the 90:10 and 50:50 mixtures of biosensor:MO cultures after exposure to ethylene (p < 0.0001), while the 10:90 mixture was less effective, and the signal from this mixture was not statistically significant (P > 0.05) (Figure 9). No significant fluorescence was seen in a 50:50 co-culture lacking ethylene, but interestingly, the biosensor culture alone with ethylene added did give a higher fluorescence in this experiment than the negative control with no biosensor cells (albeit at p > 0.05); this indicates that the mc^2^-155 host may have low levels of endogenous alkene MO activity, consistent with earlier work ^65^. The results confirm that the initial epoxide biosensor can be converted into an alkene biosensor by adding a MO enzyme to the system, and that a relatively small amount of MO activity is sufficient to give a strong biosensor response to ethylene.

### The co-culture biosensor can detect ethylene emission from fruit

As a test of the ‘real world’ functionality of the co-culture biosensor, an experiment was devised to determine whether the system could differentiate between two functionally different kinds of fresh produce, i.e. one which does not emit ethylene (carrots), and one which does emit ethylene (bananas) (Figure 10A). The co-culture biosensor was capable of distinguishing between these, and in the presence of banana, sufficient green fluorescence was obtained to be visible by eye (Figure 10B). While the fluorescent signal from the banana emissions was much weaker than the fluorescence from biosensor cultures exposed to pure ethylene oxide (e.g. Figure 10C), this experiment provides proof-of-principle that the co-culture biosensor has sufficient sensitivity to detect ethylene in a real-world context and is sufficiently selective such that other volatile emissions from non-ethylene producing items (e.g. carrots) will not interfere.

## Conclusions

Ethylene-utilising bacteria were first reported nearly 50 years ago ^66^, but subsequent progress on understanding their genetics, biochemistry and metabolism has been slow. Notably, most of the steps in the ethylene catabolic pathway and the associated CoM biosynthetic pathway are still unknown. Here, we have made an important advance via pinpointing the inducer of the ethylene oxidation pathway and by characterising the regulatory genes and promoter that respond to this chemical signal. Epoxides have been previously predicted to be the inducers of various alkene catabolic systems ^28–30^, but our work here is the first to rigorously confirm this. Due to the high homology of the NBB4 regulators to those in other ethylene and VC-oxidising isolates, we believe that epoxide-mediated induction is common to all these bacteria.

The EtnR1-EtnR2 pair is the first known example of a biological sensing system that responds to an epoxide. This is unusual due to the high reactivity and toxicity of epoxides, and their instability in aqueous solution ^67^. It is unlikely that a further-downstream metabolite is the true inducer, since the biosensor strain mc^2^-155 (pUS301-EtnR12P) lacks any of the ethylene catabolic enzymes and also lacks the ability to synthesise CoM, which is required by the second enzyme in the pathway (EtnE). It is also worth noting that although EtO hydrolyses spontaneously to yield EtGL, this by-product was a very poor inducer. The control of the ethylene catabolic system resembles that of the butane oxidation system of *T. butanivora* ^23^, where the first metabolite (1-butanol) is the inducer; we hypothesise that it is easier for bacteria to evolve sensors for functionalised molecules like alcohols and epoxides compared to the unreactive hydrocarbons that are the primary substrates of these pathways. The response of the EtO biosensor for other epoxides should be investigated in future studies.

Results from the promoter truncation assay support the initial bioinformatic analysis, indicating that the ethylene catabolic genes are controlled by a σ^70^ type promoter, and that the −35 and −10 sites have been correctly identified, despite the non-standard spacing of these elements. Although the −35 motif is less common in mycobacterial promoters compared to those of other bacteria ^54^, we found that removal of the putative −35 site from the P_etn_ sequence resulted in an 80% drop in biosensor signal, suggesting this site is very important for promoter performance in this case. The truncation of the sequence upstream of the −35 site caused some reduction in biosensor activity, but this effect was not dramatic. It might be expected that the upstream deletions would have had a larger impact on activity, since both bioinformatic analyses and the results from deletion of *etnR1* indicate that EtnR1 is an activator, and such proteins typically bind upstream of the core promoter in other characterised systems ^68,69^. Further studies to characterise inducer-protein, protein-protein, and protein-promoter binding are needed to understand the EtnR1-EtnR2-P_etn_ control system.

Key performance parameters for this biosensor were derived from the Hill transformation of the dataset, which showed a good fit (R^2^=0.9486). A Hill coefficient reflects biosensor sensitivity, where a value greater than one indicates a steeper linear response of the biosensor. The sensitivity (i.e. Hill coefficient) of pUS301-EtnR12P when induced with EtO (1.22) is broadly comparable to that of P_BMO_ (0.78-2.54 with various inducers) ^23^ and to several other common systems induced by small molecules (AraC = 1.3, CdaR =1.0) ^5^, but is lower than for BTEX biosensors such as pGLTUR (1.2−3.8 with various inducers) ^70^. The maximum fluorescence levels and linear range of detection of the EtO biosensor were also comparable to that of P_BMO_.

The co-culture assay demonstrated that the epoxide biosensor could successfully be modified to sense ethylene. Despite the complexities and limitations of this experiment (most notably, the need for the epoxide to exit one cell and enter another) the ethylene-sensing co-culture actually produced a higher fluorescent response than the epoxide-sensing single culture. This could be due to continuous production of EtO in the co-culture system, leading to a more-sustained biosensor response. Alternatively, continuous production of smaller amounts of epoxide may minimise aqueous hydrolysis or toxic effects compared to using a single larger dose of EtO. The highest signal in the co-culture system was produced by a 90:10 initial mixture of biosensor:MO cells, indicating that the regulatory components and/or reporter genes were the limiting factor rather than the MO activity; this is promising for future developments, since MO’s can be challenging to express at high levels in heterologous hosts. While the co-culture used here provided good proof-of-principle for an ethylene biosensor, future iterations of the device would involve assembly of both the MO and regulatory genes in a single plasmid; this should enable better control and functionality of the system. Integration of the whole construct (*etnABCD-etnR1-etnR2*-P_etn_-fuGFP) into the chromosome of the mc2-155 host may be desirable to enhance its stability and eliminate the need for antibiotic selection.

At face value, using a biosensor approach for detecting EtO seems ill-advised, since EtO is a potent biocide, routinely used for sterilization of medical equipment ^71^. If the biosensor were to be deployed in sterilisation contexts, it may need to remain functional in the presence of very high epoxide levels (e.g., 300 mg/L ^72^); these are several orders of magnitude higher than the detection threshold of the biosensor determined in this study. Further work is needed to determine how our biosensor would handle high concentrations of EtO, for example, in contexts such as detection of gas leaks, and at what point the EtO toxicity to the chassis bacteria would impact the function of the system. The development of cell-free transcription/translation systems (TXTL) may offer a solution to this paradox, by providing a way to apply the components of this biosensor without using live cells. It is worth noting that the alternative application as an ethylene sensor is not confounded by the analyte toxicity since ethylene itself is not particularly toxic, and only small amounts of EtO need to be generated from ethylene to trigger the biosensor.

Sensors for EtO are a necessity in workplaces that use this toxic substance since the odour threshold (260 ppm) is much higher than the allowable exposure limits. Our whole-cell EtO biosensor is more sensitive than existing commercially-available sensors based on chemical reactions; these have detection ranges of 0-10 ppm, with a resolution of 0.1 ppm ^73^, while our sensor has a detection range of 9-2,200 ppb. Although more sensitive EtO sensors do exist in the research literature which can detect parts-per-trillion EtO, these depend on expensive large equipment, i.e. an infrared laser direct absorption spectrometer ^74^. Importantly, our EtO sensor can easily detect levels corresponding to recommended workplace exposure limits (OSHA: 1 ppm over 8 h, 5 ppm in 15 min; NIOSH: 0.1 ppm per day, 5ppm in 10 min) ^75^.

The ethylene-responsive variant of our biosensor would have applications in the bulk chemical industry (e.g., monitoring workplace exposure and detection of gas leaks), but also would have applications in agriculture and in the fresh produce supply chain. Ethylene management is critical to the control of fruit ripening, whether this relates to endogenous production of the gas by the fruit itself, exposure of the produce to exogenous ethylene, or the use of chemical regulators that mimic ethylene and/or inhibit the plant ethylene receptors ^76–78^. Despite these efforts, an inability to effectively monitor fruit ripeness throughout the production chain is a major contributing factor to over 1.3 billion tonnes of food waste generated globally every year, a loss with an equivalent retail value of over USD$1 trillion that contributes ∼8% of the global greenhouse gas emissions ^79–82^. Here, we have provided proof-of-principle that our co-culture biosensor can detect ethylene emissions from bananas. Further development of this biosensor will contribute greatly to solving the intertwined problems of ripening management and minimising food waste.

## Methods

### Strains, media, solutions, and reagents

Details of strains used in this study are given in Table 2. *E. coli* TOP10 cells (Invitrogen) were used as the primary hosts for plasmid construction and were cultured in Lysogeny Broth (LB) medium ^83^. *M. smegmatis* mc^2^-155 cells ^84^ were used as the biosensor chassis; these were cultured in LB for routine purposes or Minimal Salts Medium (MSM) for fluorescence experiments ^85^. A detailed guide to preparing the MSM medium is provided in Table S1 In the Supporting Information. For mc^2^-155 cultures grown in MSM, 10 mM glucose was provided as the carbon and energy source, and 0.05% (v/v) Tween 80 was added to minimise clumping. Both *E. coli* and *M. smegmatis* cultures were grown aerobically at 37°C, with broths shaken at 200 rpm. Kanamycin was added at 50 µg/mL for *E. coli* and 20 µg/mL for *M. smegmatis*. Antibiotics, glucose, Tween80, cumate, and the trace metals solution for MSM were all added from filter-sterilized stocks after autoclaving; all these components were made as aqueous stock solutions except cumate which was dissolved in 100% ethanol. Agar was added to media where needed at 15 g/L. All molecular biology enzymes (BsaI-HFv2, T4 DNA ligase, Q5 polymerase) were supplied by NEB. Synthetic DNA fragments and oligonucleotides were supplied by either IDT or Twist Bioscience. Ethylene (99.5%), ethylene oxide (500 mg/mL in DMSO), ethylene glycol (99.8%), DMSO (99.5%), and cumic acid (98%) were from Sigma Aldrich.

**Table 2.** Plasmids and strains used in this study.

| Biological Material | Description | Genotype | Reference |
| --- | --- | --- | --- |
| *Escherichia coli* TOP10 | General purpose cloning host | F- *mcrA* $\Delta$(*mrr-hsdRMS-mcrBC*) ϕ80 *lacZ*ΔM15Δ *lacX74 recA1 ara*Δ139Δ(*ara-leu*)7697 *galU galK rpsL* (StrR) *endA1 nupG* | https://ecoliwiki.org/colipedia/index.php/TOP10 |
| *Mycobacterium smegmatis* mc<sup>2</sup>-155 | Alternative expression host | Electrocompetent, Km<sup>S</sup> | 84 |
| pUS250-fuGFP | Broad host range cumate-inducible GFP expression vector | 4750 bp, pBBR *oriV* and *mob*, *oriT*, Km<sup>R</sup>, *cymR*, *fuGFP* | https://www.addgene.org/198322/ |
| pUS116 | Acetamide-inducible mycobacterium shuttle vector | 6624 bp, *colE1 ori* (pUC), *oriM* (pAL5000), Km<sup>R</sup>, *amiC*, *amiA*, *amiD*, *amiS* | https://www.addgene.org/84578/ |
| pUS116-etnABCD | Acetamide-inducible<br>expression of<br>ethene<br>monooxygenase | 11068 bp, *colE1*<br>*ori* (pUC), *oriM*<br>(pAL5000), Km<sup>R</sup>,<br>*amiC*, *amiA*,<br>*amiD*, *amiS*,<br>*etnABCD* | 40 |
| pUS301 | Cumate-inducible<br>mycobacterium<br>shuttle vector | 5763 bp, *colE1* *ori*<br>(pUC), *oriM*<br>(pAL5000), Km<sup>R</sup>,<br>*cymR*, *fuGFP* | This study |
| pUS301-EtnR12P | Complete<br>biosensor<br>plasmid | 8363 bp, *colE1* *ori*<br>(pUC), *oriM*<br>(pAL5000), Km<sup>R</sup>,<br>*cymR*, *etnR1*,<br>*etnR2*, P<sub>etn</sub>,<br>*fuGFP* | This study |
| pUS301-EtnR1P | Biosensor<br>plasmid lacking<br>*etnR2* | 7766 bp, *colE1* *ori*<br>(pUC), *oriM*<br>(pAL5000), Km<sup>R</sup>,<br>*cymR*, *etnR1*,<br>P<sub>etn</sub>, *fuGFP* | This study |
| pUS301-EtnR2P | Biosensor<br>plasmid lacking<br>*etnR1* | 6620 bp, *colE1* *ori*<br>(pUC), *oriM*<br>(pAL5000), Km<sup>R</sup>,<br>*cymR*, *etnR2*,<br>P<sub>etn</sub>, *fuGFP* | This study |
| pUS301-EtnR12PΔ20 | Biosensor<br>plasmid with 20<br>bp deletion in<br>P<sub>etn</sub> | 6600 bp, *colE1* *ori*<br>(pUC), *oriM*<br>(pAL5000), Km<sup>R</sup>,<br>*cymR*, *etnR1*,<br>*etnR2*, P<sub>etn</sub>Δ20,<br>*fuGFP* | This study |
| pUS301-EtnR12PΔ60 | Biosensor<br>plasmid with 60<br>bp deletion in<br>P<sub>etn</sub> | 6560 bp, *colE1* *ori*<br>(pUC), *oriM*<br>(pAL5000), Km<sup>R</sup>,<br>*cymR*, *etnR1*,<br>*etnR2*, P<sub>etn</sub>Δ60,<br>*fuGFP* | This study |
| pUS301-EtnR12PΔ80 | Biosensor plasmid with 80 bp deletion in P<sub>etn</sub> | 6540 bp, *colE1 ori* (pUC), *oriM* (pAL5000), Km<sup>R</sup>, *cymR*, *etnR1*, *etnR2*, P<sub>etn</sub>Δ80, *fuGFP* | This study |
| pUS301-EtnR12PΔ100 | Biosensor plasmid with 100 bp deletion in P<sub>etn</sub> | 6520 bp, *colE1 ori* (pUC), *oriM* (pAL5000), Km<sup>R</sup>, *cymR*, *etnR1*, *etnR2*, P<sub>etn</sub>Δ100, *fuGFP* | This study |

### Plasmid construction

Details of strains used in this study are given in Table 2. pUS116 (6.6 kb) (AddGene #84578) is a *Mycobacterium*-*E. coli* shuttle vector that contains the pUC and pAL5000 origins of replication (*oriE*, *oriM*), kanamycin resistance, and an acetamidase-inducible promoter ^86^. Plasmid pUS116-etnABCD (11 kb) contains the *etnABCD* MO from *M. chubuense* NBB4 cloned into pUS116; this was made in a previous study ^40^. pUS301 (5.8 kb) was constructed in this study by amplifying the cumate regulatory system and fuGFP gene from pUS250-fuGFP with primer pair CFM3/CFM4 and cloning the PCR product into XbaI/BamHI-digested pUS116. pUS301-EtnR12P was then made by amplifying the entire pUS301 plasmid (primers CFM202/CFM206, which added BsaI sites to the ends of the amplicon), digesting with BsaI, and ligating via Golden Gate assembly with three BsaI-digested synthetic DNA fragments containing *etnR1* (1860 bp), *etnR2* (751 bp) and P_etn_ (290 bp). pUS301-EtnR1P and pUS301-EtnR2P were constructed via amplification of different sub-sections of pUS301-EtnR12P (primers CFM261/CFM262 and CFM179/CFM260, respectively), then circularisation of the amplicons by Golden Gate reaction with BsaI and ligase. pUS301-EtnR12P promoter truncation plasmids were constructed by amplifying the plasmid to exclude P_etn_ (primers CFM206/CFM294), then ligating via Golden Gate assembly with synthetic DNA fragments containing truncated P_etn_ sequences (P_etn_-20: 270 bp, P_etn_-60: 230 bp, P_etn_-80: 210 bp, P_etn_-100: 190 bp). Details of PCR primers used in this study are provided in Table S2 in the Supporting Information. Plasmid features and sequences of pUS301, pUS301-EtnR12P and pUS116-etnABCD are provided in GenBank format in Sections S2, S3, and S4 in the Supporting Information.

### Biosensor growth, induction, and fluorescence assays

The following method was used for the initial identification of inducers, characterising the response to different EtO concentrations, testing the requirement for etnR1 and etnR2, and testing truncated versions of the P_etn_ promoter. Cells of *M. smegmatis* were transformed with the biosensor plasmid (pUS301-EtnR12P or pUS301-EtnR1P or pUS301-EtnR2Pvia electroporation (2.5 kV, 800 Ω, 25 μF, 2 mm cuvette gap). After 4 days incubation, a loopful of growth (several colonies) was transferred into 5 mL LB+Km+Tween, grown to saturation (48 hours), then cells were collected by centrifugation for (15,000 *g*, 10 min). The cells were resuspended in 5 mL of MSM+Km+Tween+glucose broth, then a subsample of the cell suspension was inoculated into 50 mL of the same medium to give an initial OD_600_ of 0.02. Cultures were grown to OD_600_ = 0.25-0.50 (approx. 18-20 h), then 3 mL of culture was added to a sterile 26 mL glass vial. Cumate was added to 100 µM, then vials were crimp-sealed with Teflon-faced butyl rubber stoppers. Ethylene was added as a neat gas to give 10% v/v in the headspace, while ethylene oxide and ethylene glycol were added as a 100 μM stock solutions in DMSO to give 1 µM in the liquid phase (3 μL stock solution per 3 mL of culture). Controls lacked either cumate or the test inducers; in the latter case, DMSO only was added. The vials were incubated horizontally with shaking to ensure good gas exchange. Sampling of cells was done 16-20 h post-induction via extracting 200 µL of culture with a syringe and transferring the sample to a 96-well plate (polystyrene, flat-bottom, black-edged, Greiner #655090). Fluorescence was measured in a plate reader (ClarioStar Plus, BMG Labtech) at 470 nm excitation and 515 nm emission, in both cases with a bandwidth of 20 nm, with the gain set to 1000. The culture turbidity (OD_600_) was measured at the same time, and the RFU calculated as fluorescence/OD_600_. For the EtO concentration curve, samples were collected 10 hours post-induction with cumate and inducer.

For the time course assay, the above method was modified by scaling the system up to 24 mL cultures in 120 mL crimp-sealed bottles, and by sampling at multiple time points over a 29-hour incubation. For testing the effect of pre-induction of the regulators, the original protocol above was modified by adding cumate at the point of transfer of the starter culture into the 50mL broth, and by adding EtO 16-20 h later, after the culture had reached an OD_600_ = 0.25-0.3. For the co-culture assay, the original protocol was modified by inclusion of a second culture of mc^2^-155, carrying plasmid pUS116-etnABCD; these cells were prepared similarly to the mc^2^-155(pUS301-EtnR12P) cells (i.e., freshly transformed with plasmid, pre-grown in a 5 ml broth, then transferred to a 50 ml broth), except that they were induced with acetamide (1%) instead of cumate at the point of transfer into the 50 ml broth. In this experiment, the pre-induction method was used for the mc^2^-155(pUS301-EtnR12P) culture, so it was also induced with cumate at the point of transfer into the larger broth. After the respective inducer additions, both cultures were grown until OD600 = 0.25-0.50 (∼18 hours), then the cultures were mixed at different volumetric ratios (50:50, 90:10, 10:90) in the 26 ml test vials to give a total culture volume of 3 mL. After crimp-sealing, ethylene was added to the headspace (2% v/v), the cultures incubated 16 hours with shaking, then fluorescence and OD_600_ were measured in the plate reader as described above, and RFU calculated. GraphPad Prism was used to fit the Hill equation to the data set and calculate the Hill coefficient and K_m_ values (Sigmoidal, 4PL, X is concentration, nonlinear regression). Hill equation: 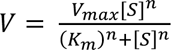, where V is the response, S is the substrate, K_m_ is the substrate concentration that gives half the maximum response, and n is the Hill coefficient.

### Detection of ethylene emission from fruit

Cultures of mc^2^-155(pUS301-EtnR12P) and mc^2^-155(pUS116-etnABCD) were separately grown in 50 mL MSM+Km+Tween+glucose broths, induced with cumate or acetamide, respectively, and harevested at mid-exponential phase, as described above. The cells were centrifuged and resuspended to OD_600_ = 5 in the same medium, then mixed at a ratio of 2:1 (biosensor cells: epoxidating cells) in 3 ml total volume and added into 16 mL glass vials with no cap. Fresh produce items, either carrot or banana, were sliced into 3 mm thick discs which were then further chopped in half, then 10 g of material was added into separate 100 mL Schott bottles. The vials containing the biosensor co-culture were suspended with string inside each bottle, then the bottles were tightly capped, and incubated at room temperature for 3 days with shaking (150 rpm). The fluorescence of the cell suspensions was then visualised qualitatively under long-wave UV light (405 nm emission maximum) using hand-held torch.

## Author Contributions

CM, NC, and CS designed the study. CM and NY performed lab work, under the supervision of NC and CS. CM prepared the first draft of the manuscript, which was then edited by NC and CS.

## Supporting Information

Section S1. Calculation of ethylene oxide (EtO) in aqueous phase using Henry’s constant

Figure S1. Growth of mc^2^-155(pUS301-EtnR12P) cultures after induction

Table S1. Preparation of minimal salts medium (MSM)

Table S2. PCR primers used in this study

Section S2. Sequence and features of plasmid pUS301-EtnR12P

Section S3. Sequence and features of plasmid pUS301-EtnR12P

Section S4. Sequence and features of plasmid pUS116-etnABCD

## Acknowledgements

CM was supported during her PhD candidature by a Research Training Program scholarship from the Australian Government, and by a top-up scholarship and operating costs budget from a CSIRO Synthetic Biology Future Science Platform PhD award. We acknowledge the University of Sydney 2016 iGEM team ‘FRESH’ (Amanda Chen, Sing-Young Chen, Sholto Douglas, Liam Ferguson, Wunna Kyaw, Claudia Moratti, Orion Tong, Alma Wu) for their contributions to preliminary work on this project.

## Development of a whole-cell biosensor for ethylene oxide and ethylene

### Supporting information

**Section S1.**
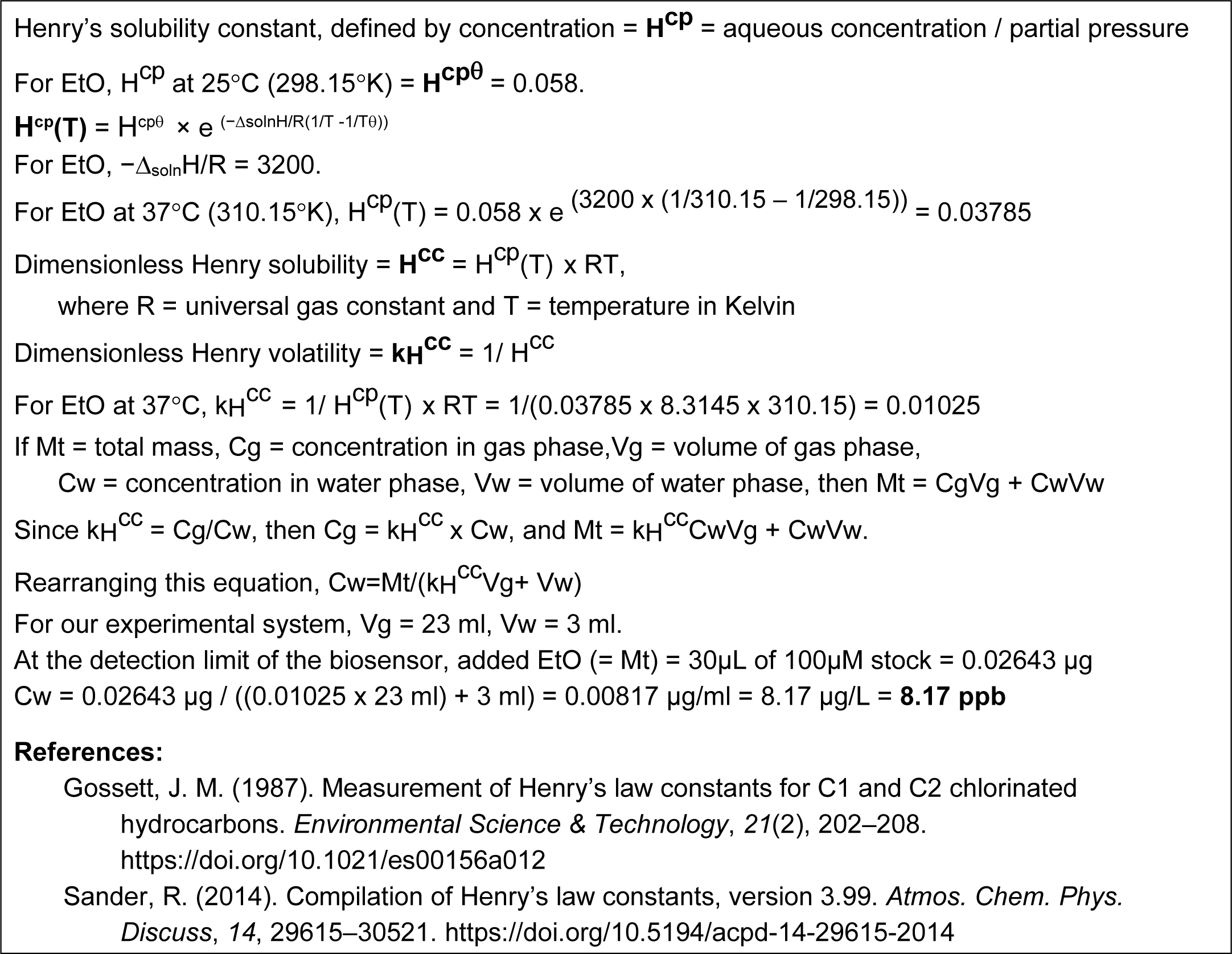
Calculation of ethylene oxide (EtO) in aqueous phase using Henry’s constant.

**Figure S1.**
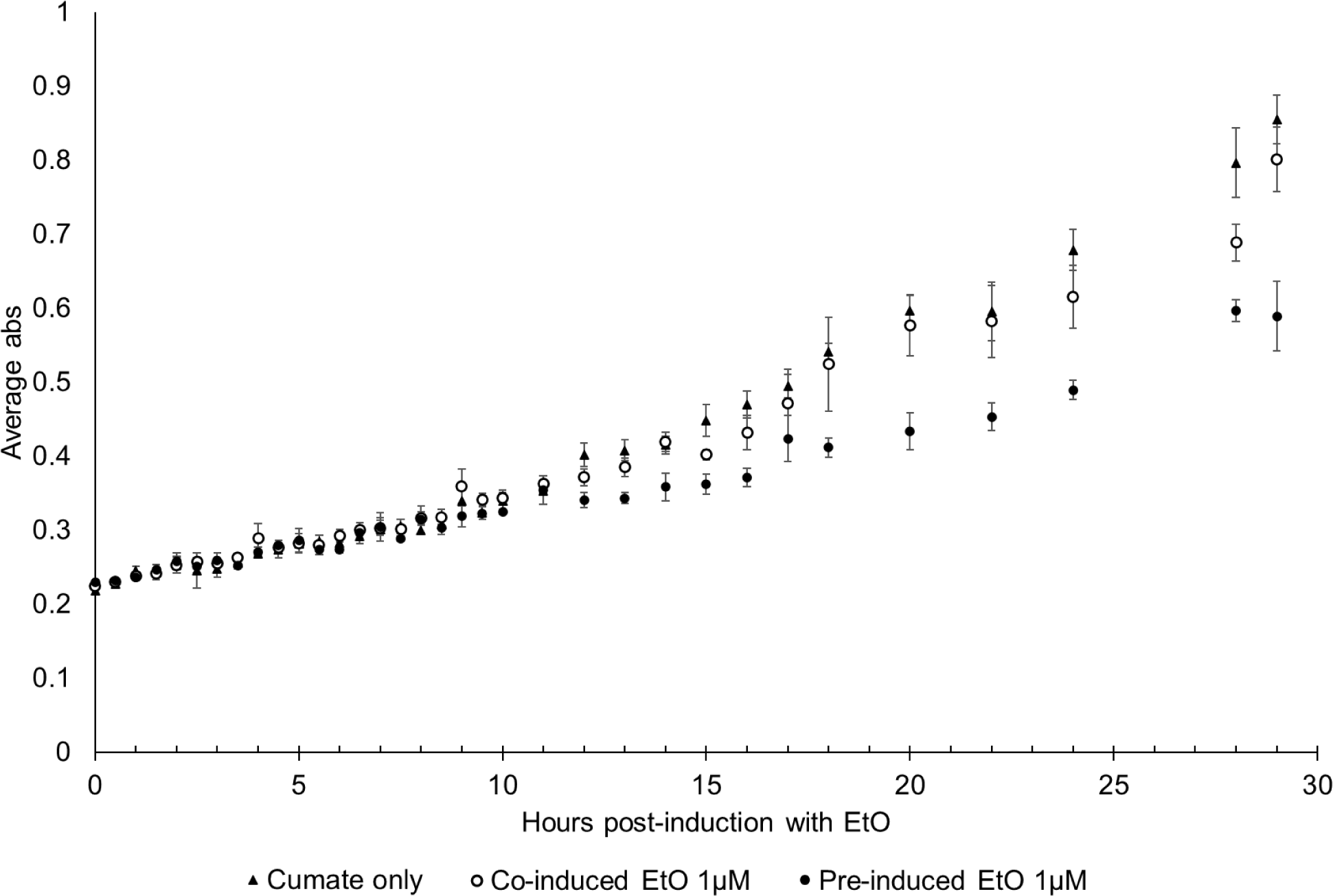
Growth of mc^2^-155(pUS301EtnR1R2P) cultures after induction. Growth in this experiment was measured as OD600. Data are the means of three replicates, and error bars show the standard deviation.

**Table S1.** Preparation of minimal salts medium (MSM) The MSM bulk medium is prepared first, adjusted to pH 7.0 with H_2_SO_4_, and sterilised by autoclaving, then trace elements solution and carbon sources are added after cooling. Trace elements solution is adjusted to pH 6.5 with H_2_SO_4_, sterilised by filtration through a 0.22 µm filter, then stored at 4°C, wrapped in foil.

| Bulk medium components | Amount added per litre |
| --- | --- |
| K<sub>2</sub>HPO<sub>4</sub> | 2.27 g |
| KH<sub>2</sub>PO<sub>4</sub> | 0.95 g |
| (NH<sub>4</sub>)<sub>2</sub>SO<sub>4</sub> | 0.67 g |
| Trace elements solution | 2 ml |
| Trace elements components | Amount added per litre |
| Na<sub>2</sub>EDTA.2H<sub>2</sub>O | 6.37 g |
| ZnSO<sub>4</sub>.7H<sub>2</sub>O | 1.0 g |
| CaCl<sub>2</sub>.2H<sub>2</sub>O | 0.5 g |
| FeSO<sub>4</sub>.7H<sub>2</sub>O | 2.5 g |
| NaMoO<sub>4</sub>.2H<sub>2</sub>O | 0.1 g |
| CuSO<sub>4</sub>.5H<sub>2</sub>O | 0.1 g |
| CoCl<sub>2</sub>.6H<sub>2</sub>O | 0.2 g |
| MnSO<sub>4</sub>.H<sub>2</sub>O | 0.52 g |
| MgSO<sub>4</sub>.7H<sub>2</sub>O | 60.0 g |

**Table S2.**
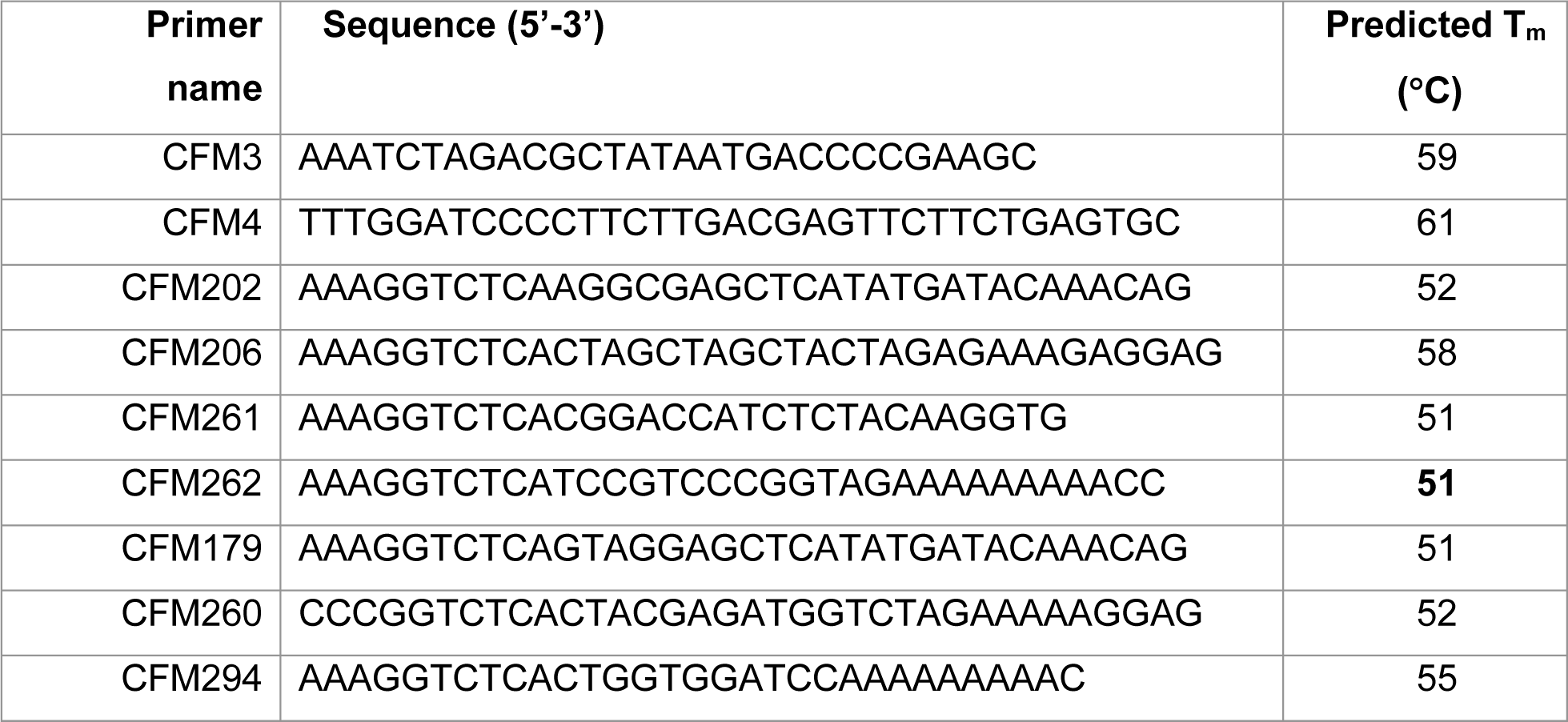
PCR primers used in this study.

**Section S2.**
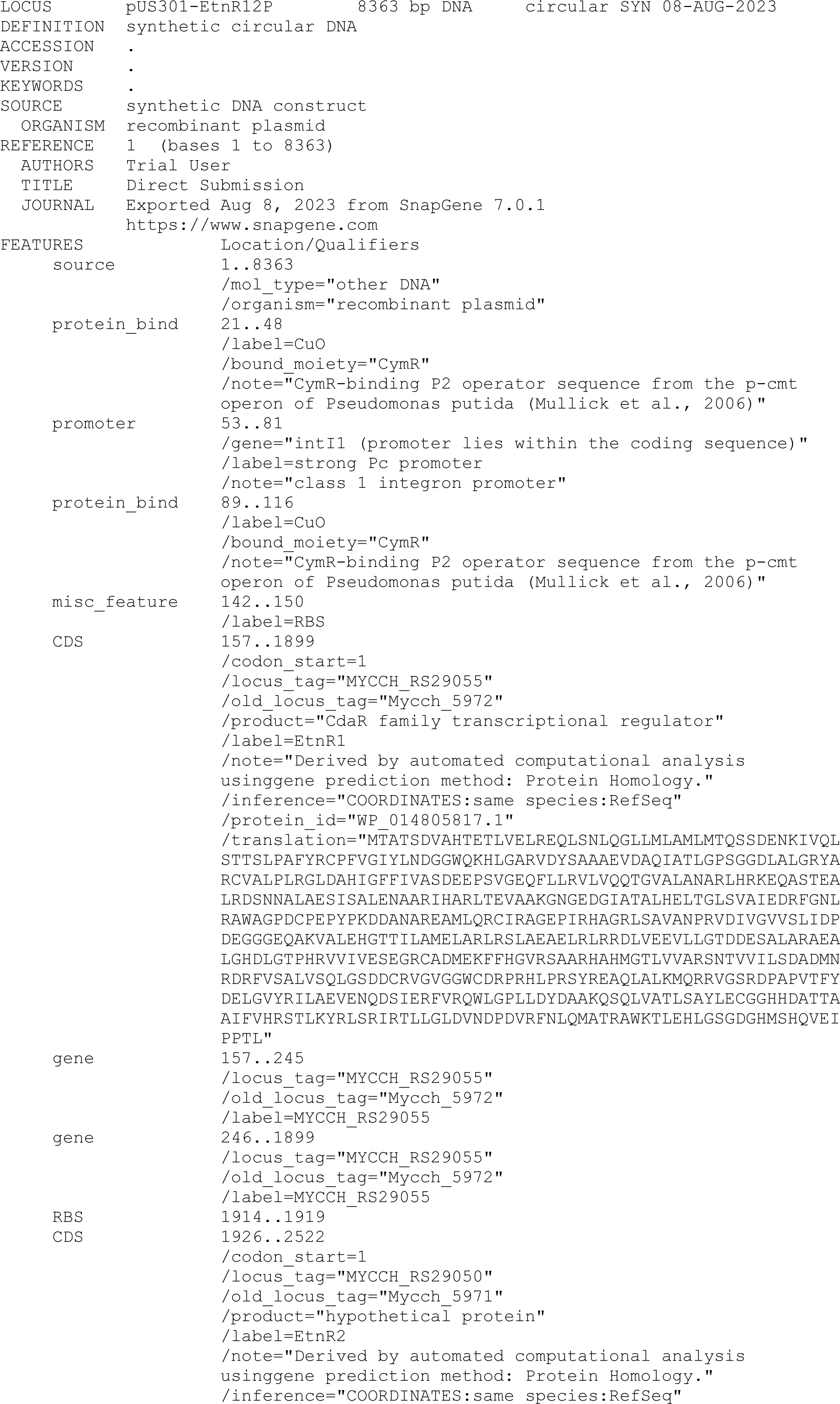
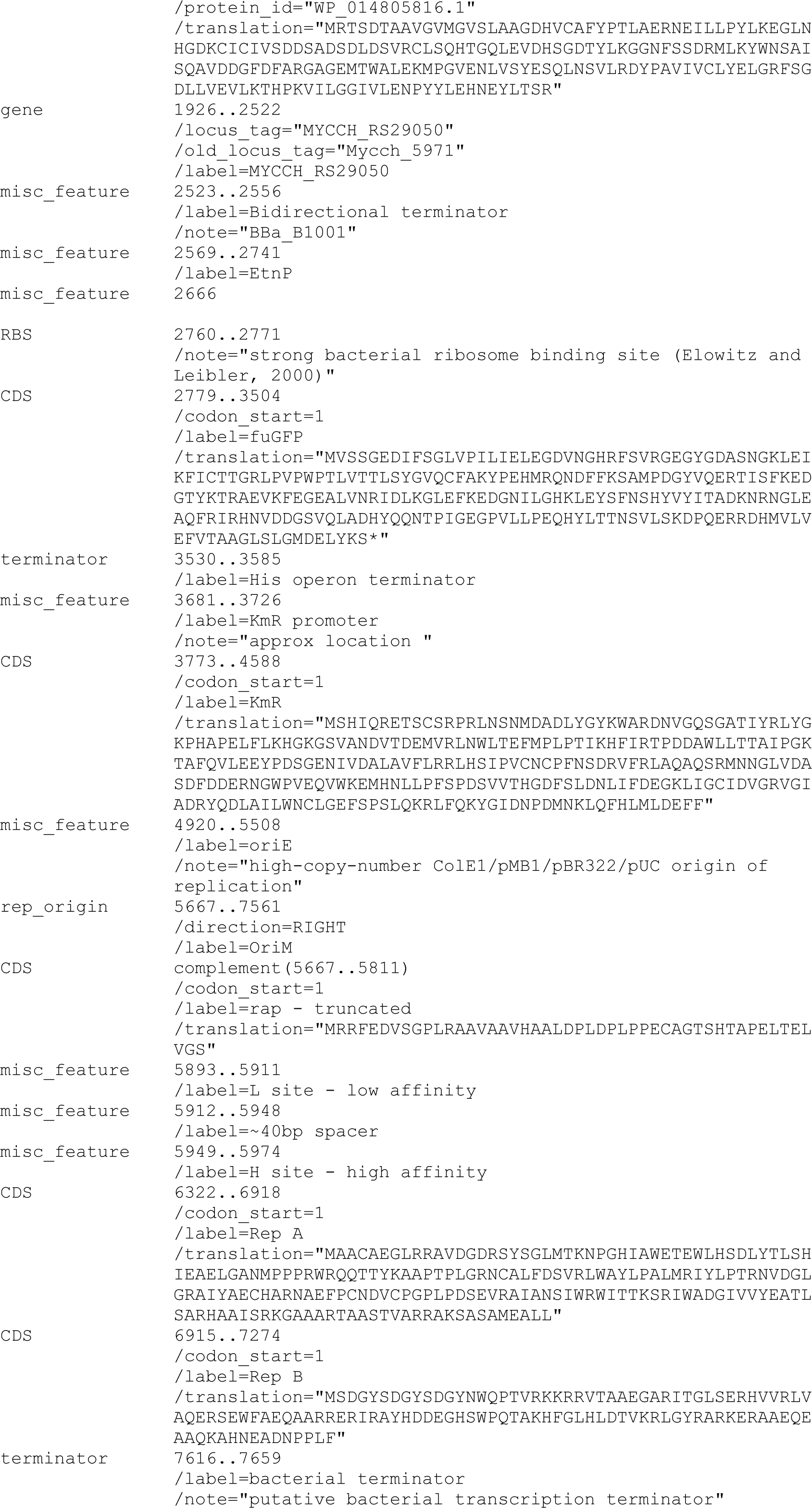
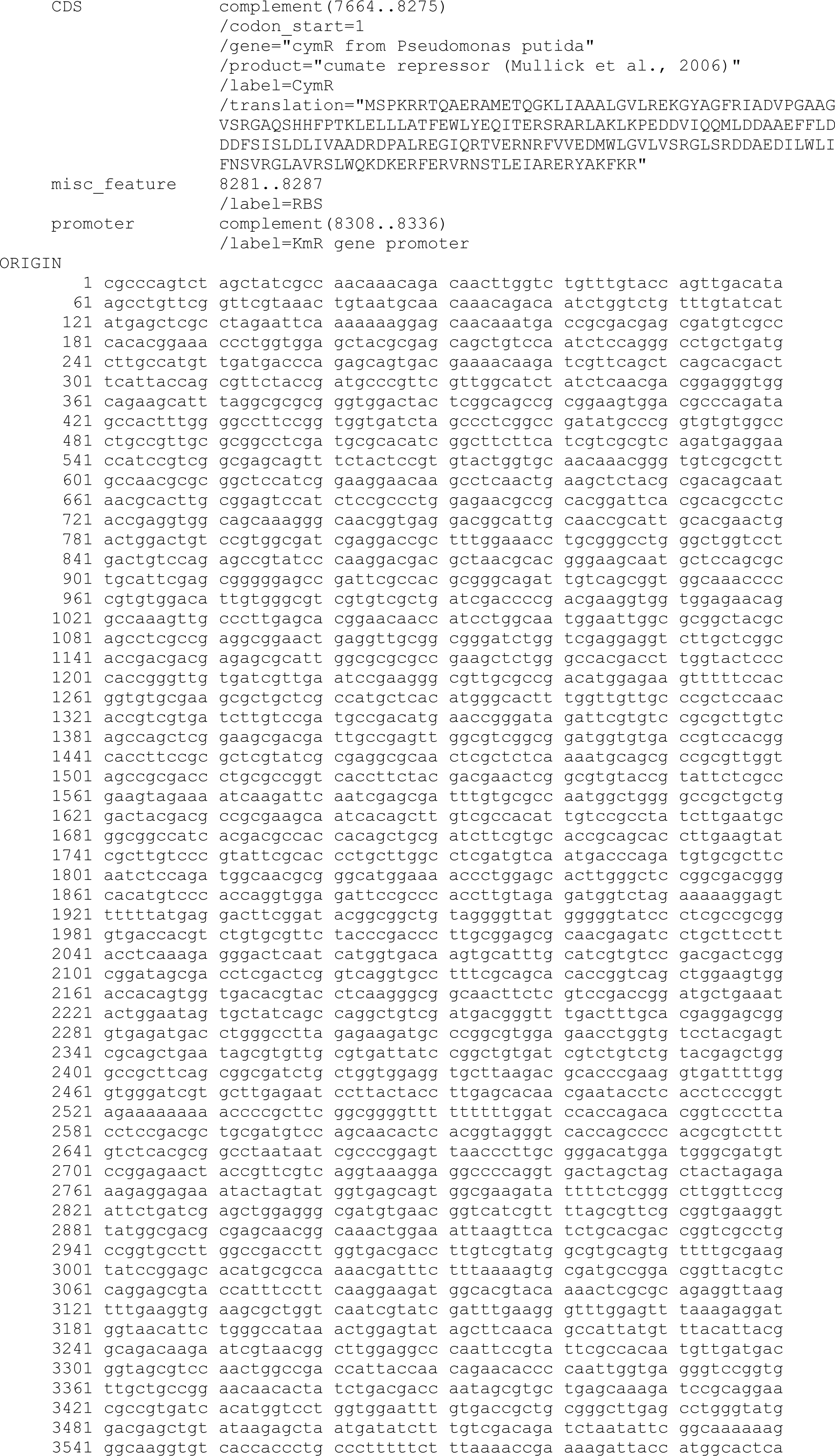
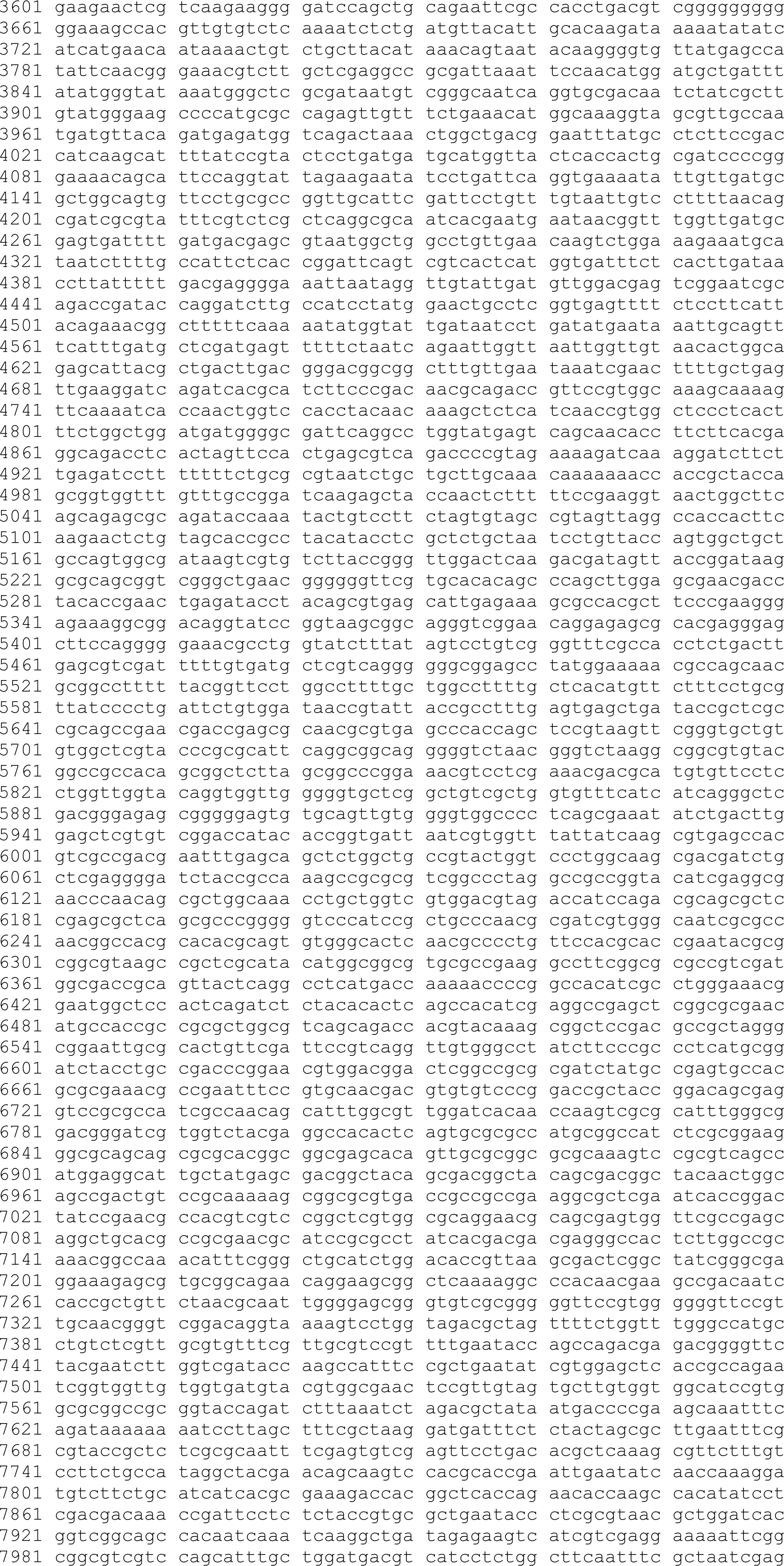
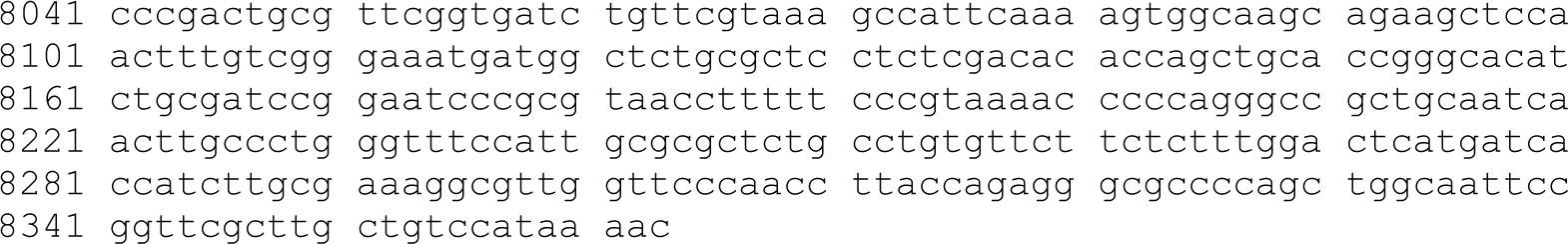
Sequence and features of plasmid pUS301-EtnR1R2P.

**Section S3.**
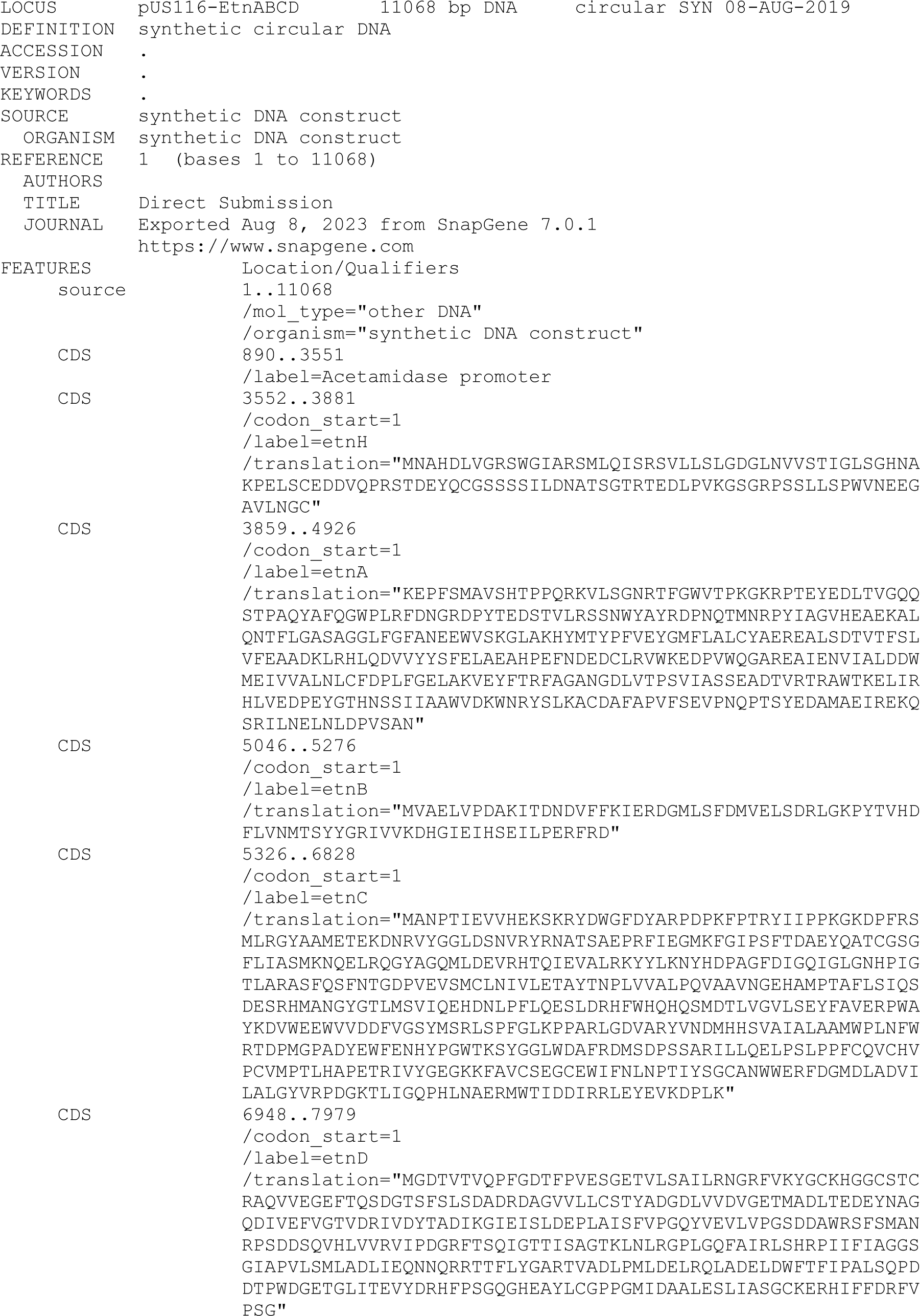
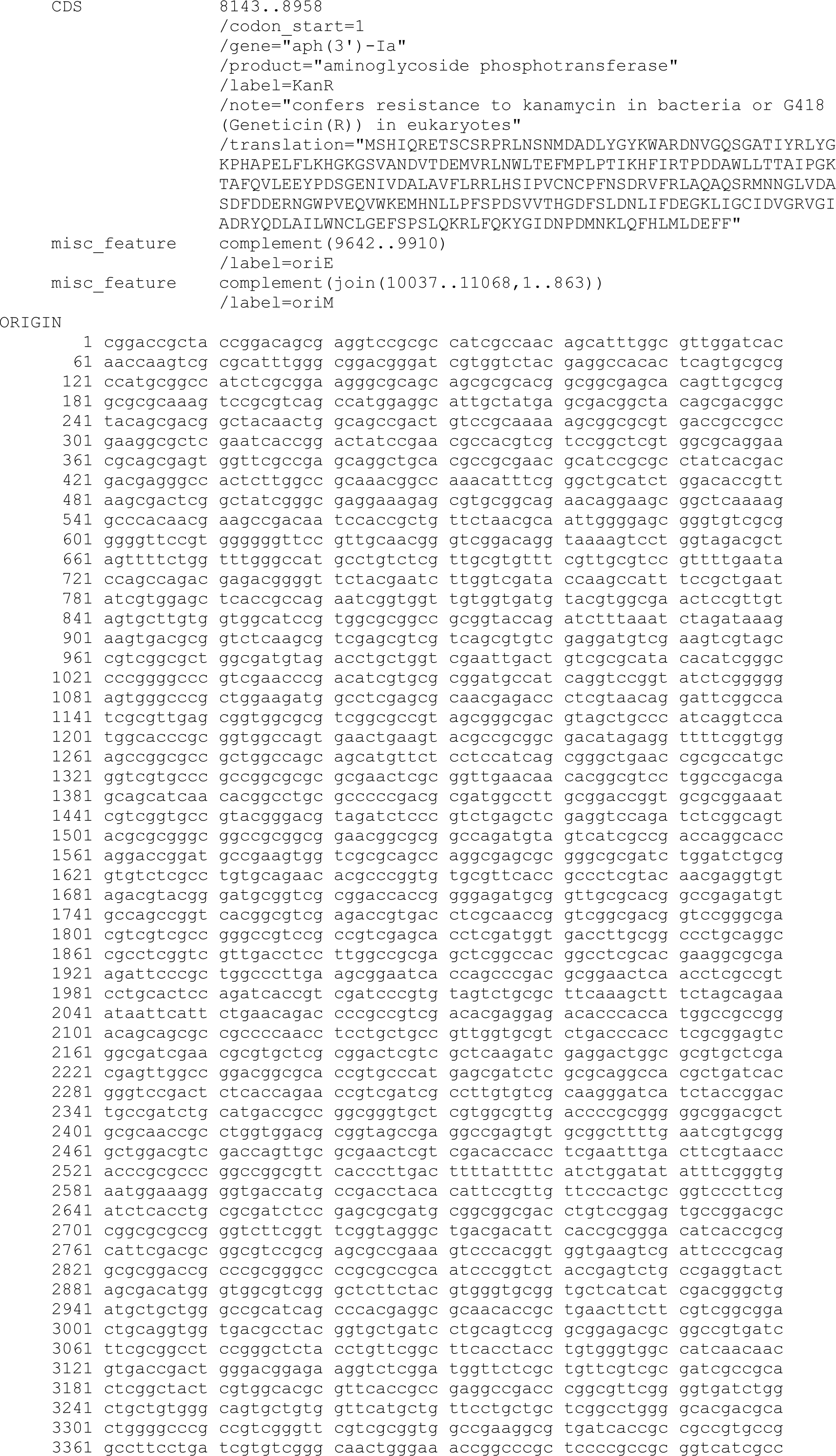
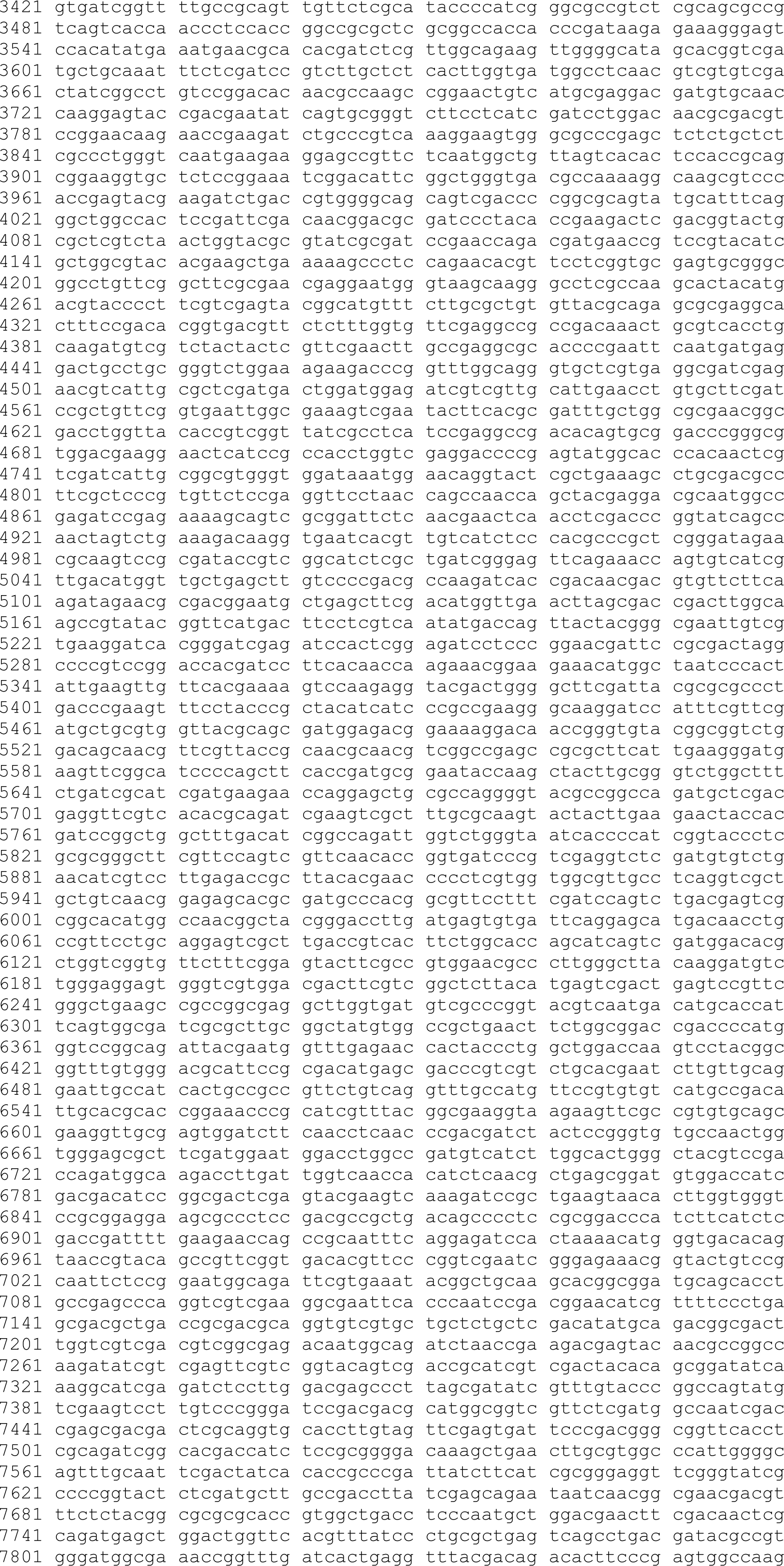
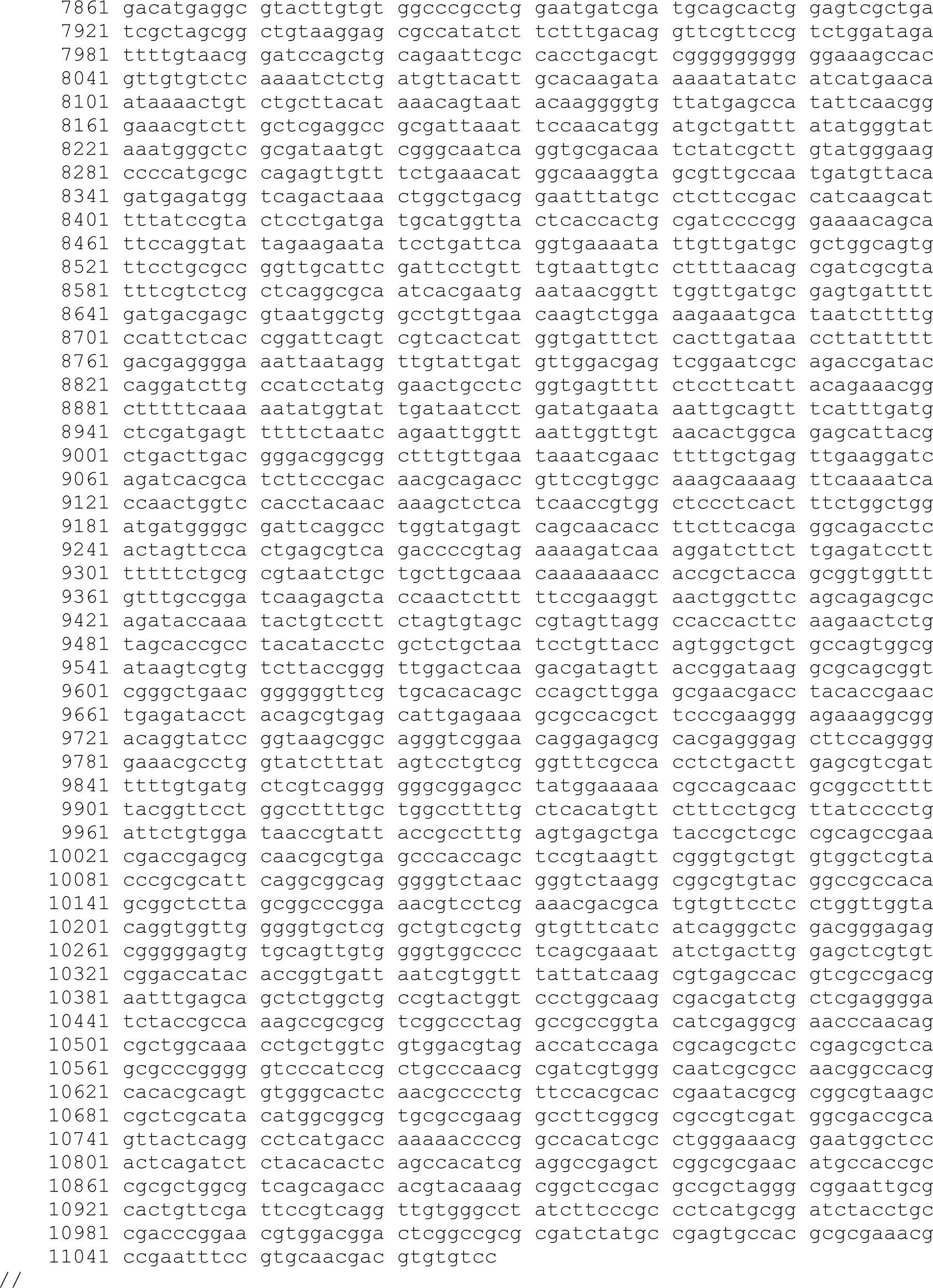
Sequence and features of plasmid pUS116-EtnABCD.

